# Tumor growth of neurofibromin-deficient cells is driven by decreased respiration and hampered by NAD^+^ and SIRT3

**DOI:** 10.1101/2021.05.29.446262

**Authors:** Ionica Masgras, Giuseppe Cannino, Francesco Ciscato, Carlos Sanchez-Martin, Marco Pizzi, Alessio Menga, Alessandra Castegna, Andrea Rasola

**Affiliations:** Department of Biomedical Sciences, University of Padova, Viale G. Colombo 3, 35131, Padova, Italy; Institute of Neuroscience, National Research Council, Viale G. Colombo 3, 35131, Padova, Italy; General Pathology and Cytopathology Unit, Department of Medicine-DIMED, University of Padova, via Giustiniani 2, 35128, Padova, Italy; Department of Biosciences, Biotechnologies and Biopharmaceutics, University of Bari, via Orabona 4, 70125, Bari, Italy; IBIOM-CNR, Institute of Biomembranes, Bioenergetics and Molecular Biotechnologies, National Research Council, Via G.Amendola 122/O, 70126, Bari, Italy

## Abstract

Neurofibromin loss drives neoplastic growth and a rewiring of mitochondrial metabolism. Here, we report that neurofibromin ablation dampens expression and activity of NADH dehydrogenase, the respiratory chain complex I, in an ERK-dependent fashion. This provides cells with resistance to pro-oxidants targeting complex I and decreases both respiration and intracellular NAD^+^. Expression of the alternative NADH dehydrogenase NDI1 raises NAD^+^/NADH ratio, enhances the activity of the mitochondrial NAD^+^-dependent deacetylase SIRT3 and interferes with tumorigenicity in neurofibromin-deficient cells. This anti-neoplastic effect is mimicked both *in vitro* and *in vivo* by administration of NAD^+^ precursors or by rising expression of the NAD^+^ deacetylase SIRT3, and is synergistic with ablation of the mitochondrial chaperone TRAP1, which augments succinate dehydrogenase activity further contributing to block pro-neoplastic metabolic changes of these cells. These findings shed light on chemotherapeutic resistance and on bioenergetic adaptations of tumors lacking neurofibromin, linking complex I inhibition to mitochondrial NAD^+^/NADH unbalance and SIRT3 inhibition, as well as to down-regulation of succinate dehydrogenase. This metabolic rewiring could unveil attractive therapeutic targets for neoplasms related to neurofibromin loss.

## INTRODUCTION

Metabolic changes confer tumor cells the capability to adapt to heterogeneous and changing environmental conditions, promoting neoplastic progression. Bioenergetic rewiring results from a complex interplay of many factors, both intrinsic to the tumor type and enacted by interactions with heterotypic cellular and matrix components (Kim and DeBerardinis, 2019; Vander Heiden and DeBerardinis, 2017). Mitochondria sense and integrate these fluctuating signals and orchestrate adaptive metabolic responses necessary to provide building blocks to the rapidly proliferating cancer cells, while shielding them from ROS damage and hypoxic conditions in a context of oncogenic signaling dysregulation (Cannino et al., 2018; Vyas et al., 2016; Zong et al., 2016).

In neoplastic models characterized by hyperactivation of Ras-driven transduction pathways, cells undergo profound metabolic alterations characterized by down-regulation of oxidative phosphorylation (OXPHOS), anaplerotic activation of glutamine utilization as carbon source for tricarboxylic acid (TCA) cycle and induction of the (pseudo)hypoxic transcriptional program coordinated by HIF1 (Cannino et al., 2018; Kawada et al., 2017; Kim and DeBerardinis, 2019; White, 2013). Aberrant induction of Ras signaling is found in Neurofibromatosis type 1 (NF1), a tumor-predisposing genetic disorder caused by loss of function mutations of the NF1 gene encoding the Ras-GAP (GTPase-activating protein) neurofibromin (Yap et al., 2014). Deregulation of Ras/MEK/ERK transduction cascade is mandatory for tumorigenicity of Nf1^-/-^ cells (Ratner and Miller, 2015). NF1 patients are prone to develop diverse tumor types and are hallmarked by the onset of neurofibromas, benign neoplasms that affect Schwann cells and can evolve to extremely aggressive MPNSTs (malignant peripheral nerve sheath tumors) (Ratner and Miller, 2015). Metabolic features of neurofibromin-deficient cells as well as their regulatory mechanisms have been poorly investigated, even though it is known that transformation of neurofibromas into MPNSTs is accompanied by an increased avidity for glucose (Meany et al., 2013; Tovmassian et al., 2016), an observation that could open therapeutic possibilities for these poorly manageable sarcomas.

We have demonstrated that precise post-translational modifications (PTMs) of key regulatory molecules induce a fine metabolic tuning that influences neoplastic progression in cells lacking neurofibromin. Activity of succinate dehydrogenase (SDH) is decreased by interaction with the mitochondrial chaperone TRAP1 (Sciacovelli et al., 2013), and its inhibitory function is further enhanced by ERK-dependent phosphorylation. The consequent increase in intracellular succinate levels stabilizes HIF1α and installs a pseudohypoxic phenotype required for NF1-related tumor growth (Masgras et al., 2017a). Additional PTMs are provided by sirtuins, a class of protein deacylases the catalytic activity of which requires the metabolic cofactor nicotinamide adenine dinucleotide (NAD^+^) (Chalkiadaki and Guarente, 2015; Houtkooper et al., 2012). The NAD^+^/NADH pair participates in a variety of metabolic reactions that entail an electron exchange, shaping glycolysis, tricarboxylic acid cycle, oxidative phosphorylation and fatty acid oxidation; moreover, NAD^+^ is the precursor for NADP^+^ and NADPH, which display important anti-oxidant and biosynthetic functions (Katsyuba and Auwerx, 2017). Hence, NAD^+^-dependence make sirtuins cellular metabolic sensors, and in neoplastic cells they couple the bioenergetic status with a signaling output affecting tumorigenicity (Anderson et al., 2017).

In mitochondria, a fundamental role in bioenergetic regulation is accomplished by SIRT3, which activates several enzymes of TCA cycle, OXPHOS and fatty acid oxidation, thus increasing their functional coordination for ATP synthesis (Torrens-Mas et al., 2017; Zhu et al., 2019). SIRT3 roles in the neoplastic process are controversial (Carafa et al., 2019): its expression is upregulated in several types of cancers, where it has been referred to as an oncogene preventing apoptosis and promoting cell proliferation (Zhao et al., 2019). Conversely, SIRT3 ablation favors tumor growth in mice, and a tumor suppressor role of SIRT3 has been reported in breast cancer, hepatocellular carcinoma, metastatic ovarian cancer and B-cell malignancies (Torrens-Mas et al., 2017; Xiong et al., 2016). This dichotomous role in cancer progression could depend on the type, stage and microenvironment of the tumor, but also on NAD^+^ availability. Moreover, SIRT3 activates the ROS-scavenging enzyme manganese superoxide dismutase (MnSOD) and isocitrate dehydrogenase (IDH), which generates the anti-oxidant NADPH (Tao et al., 2010; Torrens-Mas et al., 2017). Therefore, in the absence of SIRT3, mitochondrial ROS levels increase, leading to multifaceted and context-dependent effects on the tumorigenic process. For instance, SIRT3 opposes the pseudo-hypoxic, pro-neoplastic phenotype mastered by HIF1α, as the increase in ROS caused by loss of SIRT3 leads to HIF1α stabilization (Finley et al., 2011a).

In this study, we report that absence of neurofibromin lowers the quantity and activity of complex I of the electron transport chain, which is the entry point for high-energy electrons from NADH into the OXPHOS (Sazanov, 2015), thus diminishing intracellular NAD^+^/NADH ratio. We provide evidence that increasing intracellular NAD^+^ and reactivating SIRT3 synergize with TRAP1 inhibition in ablating neoplastic growth of NF1-related malignancies, unveiling a potential therapeutic option grounded on dissection of their metabolic features.

## RESULTS

### Neurofibromin loss decreases protein levels and enzymatic activity of NADH dehydrogenase in an ERK1/2 dependent manner

We compared mitochondrial OXPHOS complexes in mouse embryonic fibroblasts (MEFs) derived from wild type and Nf1^-/-^ animals (Shapira et al., 2007), finding that Nf1^-/-^ MEFs had a lower expression and activity of respiratory chain complex I (*aka* NADH dehydrogenase or NADH ubiquinone oxidoreductase) than their wild-type counterparts (Figures 1A-1C). Similarly, Nf1^-/-^ MEFs exhibited a decrease of complex I expression and activity in respiratory supercomplexes (SCs) (Figure 1D) that functionally link OXPHOS complexes to funnel electron transfer along the respiratory chain (Enriquez, 2016; Lenaz et al., 2016). Expression of the GAP-related domain of neurofibromin (NF1-GRD), which reverses Ras activation in Nf1^-/-^ MEFs (Shapira et al., 2007), rescued complex I activity (Figures S1A) while the constitutively active Ras^G12D^ mutant decreased it in wild-type MEFs, mimicking the effect of neurofibromin ablation (Figures S1B).

**Figure 1.**
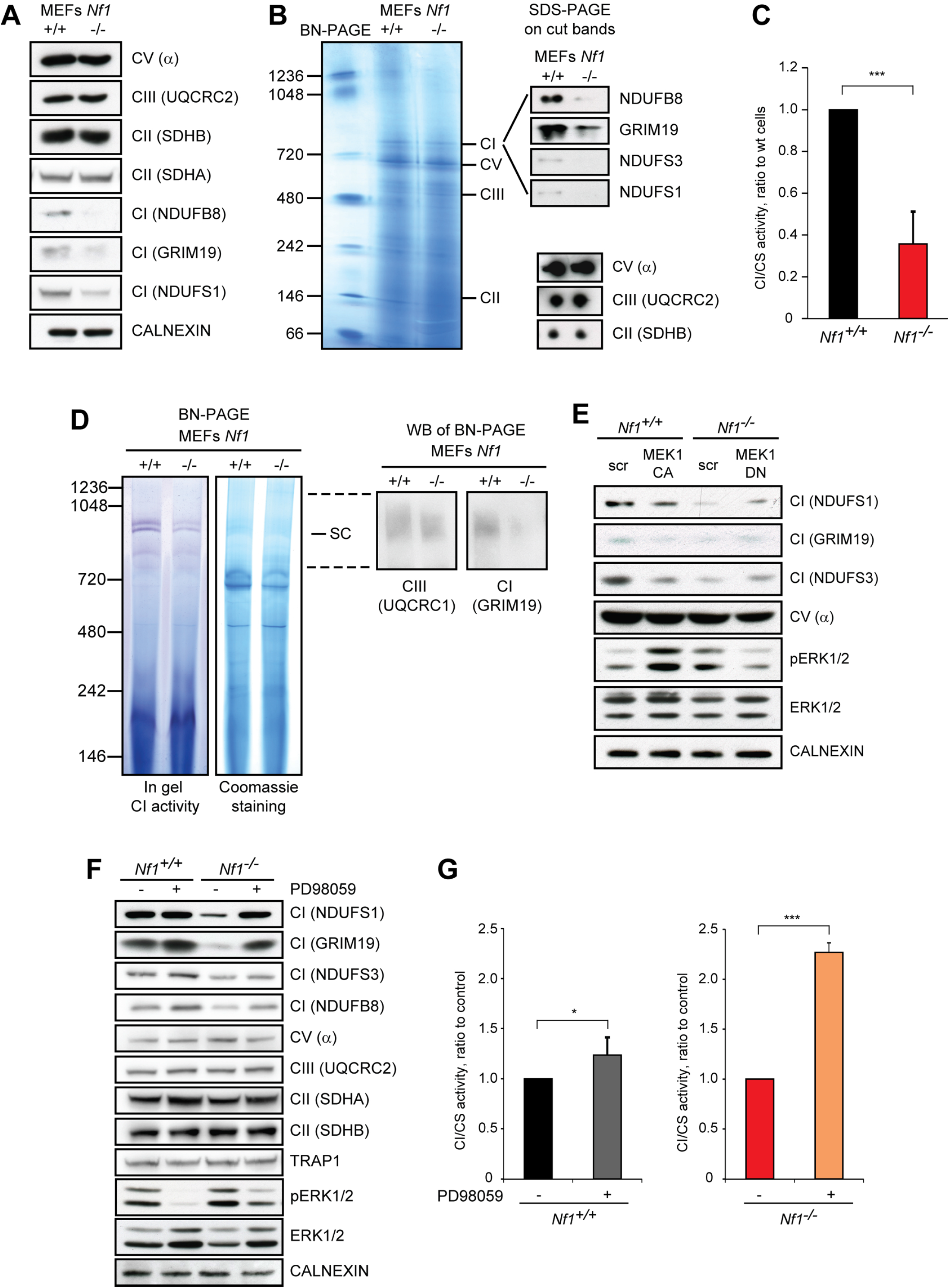
The absence of neurofibromin decreases protein levels and enzymatic activity of respiratory complex I in an ERK1/2 dependent manner. (A and B) OXPHOS protein levels were analyzed by Western immunoblot (WB, A) or blue native polyacrylamide gel electrophoresis (BN-PAGE; B). In (A), calnexin was used as a loading control. In (B), bands corresponding to different respiratory complexes were cut, run on a SDS-PAGE and probed for the expression of the complex I subunits NDUFB8, GRIM19, NDUFS1 AND NDUFS3, of the complex V α subunit, of the complex III UQCRC2 subunit and of the complex II SDHB subunit. (C) Spectrophotometric analysis of the NADH dehydrogenase activity of complex I (CI) is shown as arbitrary units and normalized for citrate synthase (CS) activity. Data are reported as mean ± SD values (n ≥ 3); ***:p<0.001; **: p<0.01; *: p<0.05 with a Student’s *t* test analysis. (D) BN-PAGE carried out on digitonized mitochondria to preserve assembled respiratory supercomplexes. Gels were either stained with Coomassie blue and transferred to PVDF membrane for protein quantification, or subjected to in gel complex I activity. UQCRC1 and GRIM19 are complex III and complex I subunits, respectively. (E and F) Western immunoblot analyses of Complex I subunits upon modulation of ERK activity through expression of the constitutively-active or dominant-negative form of the upstream MEK1 kinase (MEK1-CA and MEK1-DN, respectively; E) or by treatment with the MEK inhibitor PD98059 (40 µM, 3 days; F). NDUFS1, GRIM19, NDUFS3 and NDUFB8 were used as complex I markers, subunits α, UQCRC2 and SDHA/B as complex V, complex III and complex II markers, respectively. pERK1/2 indicates phosphorylated, active ERK1/2. TRAP1 and calnexin were used as loading controls. (G) Complex I activity with or without PD98059 treatment (40 µM, 3 days) was analyzed as in (C). All experiments in the Figure were carried out in Nf1^+/+^ and Nf1^-/-^ MEFs. See also Figure S1.

This down-regulation of complex I was elicited by Ras/MEK/ERK signaling activation, as demonstrated by expression of a constitutively active MEK1 (MEK1-CA) kinase, whereas ERK inhibition by a dominant-negative MEK1 (MEK1-DN) protein, as well as by the MEK inhibitor PD98059, enhanced both protein levels and enzymatic function of NADH dehydrogenase (Figures 1E-G and Figures S1C and S1D). MEK inhibition correlated with an increase in complex I expression also in human U87 glioblastoma cells (Figure S1E) and ipNF 04.4 plexiform neurofibroma cells (Figure S1F), both characterized by absence of neurofibromin and hence by Ras/MEK/ERK pathway induction. Protein levels of the other OXPHOS complexes were unaffected by hyperactivation of Ras/MEK/ERK signaling in all these cell models (Figures 1A, 1B and 1E-1F and Figures S1C-S1F). Neurofibromin loss did not elicit any difference in mRNA expression of complex I subunits (Figure S1G). These results connect deregulated activation of Ras/MEK/ERK signaling caused by absence of neurofibromin with complex I inhibition, in keeping with a reported role of oncogenic Ras in orchestrating the metabolic rewiring of tumor cells (White, 2013). Therefore, we investigated the possible effects of complex I inhibition on the neoplastic features of Nf1^-/-^ cells.

### Absence of neurofibromin decreases mitochondrial ROS and sensitivity to drugs targeting complex I

NADH dehydrogenase is one of the major sources of cellular ROS (Hirst and Roessler, 2016). Accordingly, we found that Nf1^+/+^ MEF cells, which display a higher complex I activity than Nf1^-/-^ MEFs (Figures 1C, 1D and 2A), have a higher level of mitochondrial ROS (Figure 2B). Several widely used chemotherapeutics act at least in part by eliciting a ROS surge in tumor cell mitochondria (Gorrini et al., 2013), and the development of compounds that prompt oxidative stress and cell death by targeting complex I could constitute an interesting anti-neoplastic strategy. One of these molecules, the gold-dithiocarbamate complex AUL12 (dibromo [ethyl-*N*-(dithiocarboxy-kS,kS′)-*N*-methylglycinate] gold(III)), initiates a composite apoptotic signaling in tumor cells following complex I inhibition and the ensuing ROS induction (Chiara et al., 2012; Nardon et al., 2015). Exposure to AUL12 inhibited complex I activity and increased mitochondrial ROS in Nf1^+/+^ MEF cells (Figures 2A and 2B), leading to oxidative stress-dependent cell death (Figure 2C). However, AUL12 could not induce either ROS surge or cell death in Nf1 knock-out cells, in which the low NADH dehydrogenase activity is not inhibited by the compound in a statistically significant manner (Figures 2A-2C). Similarly, the classical complex I inhibitor rotenone induced higher mitochondrial ROS levels and prompted more cell death in Nf1^+/+^ than in Nf1^-/-^ MEF cells (Figures 2D and 2E). Other compounds endowed with a pro-oxidant effect on mitochondria at least partially caused by inhibition of complex I, *i*.*e*. the BH3 mimetic EM20-25 and cisplatin (Ciscato et al., 2014), showed a higher toxicity on wild-type than on Nf1^-/-^ MEFs (Figures S2A and S2B). ROS scavenging protected cells from death induced by all these treatments (Figures 2C and 2E and Figures S2A and S2B). Instead, complex III inhibitor Antimycin A (AA) induced the same amount of mitochondrial ROS independently of the presence of neurofibromin (Figure S2C). Our results indicate that Ras/MEK/ERK-dependent inhibition of NADH dehydrogenase activity confers resistance to environmental conditions or pharmaceutical treatments that cause oxidative stress.

**Figure 2.**
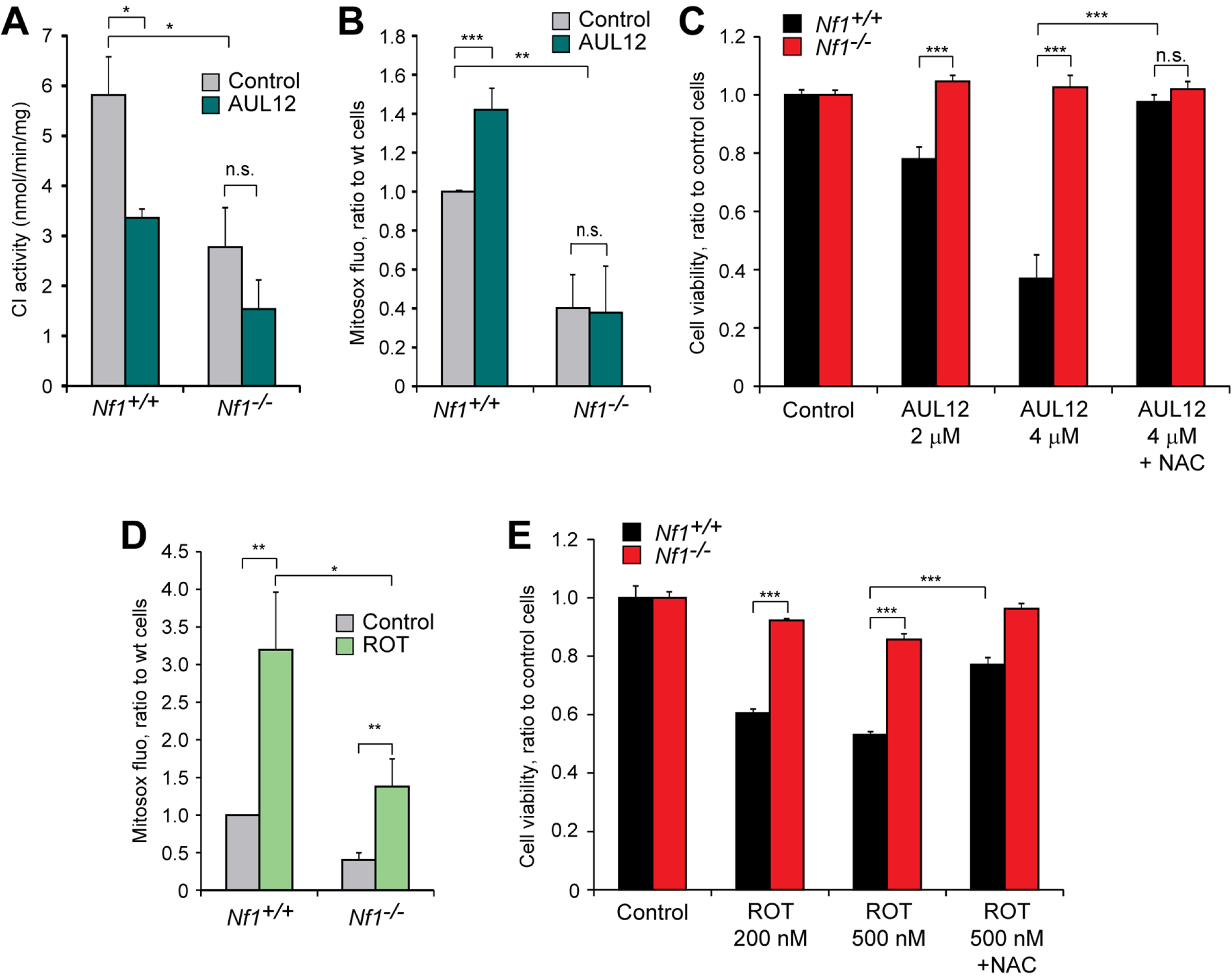
The absence of neurofibromin decreases mitochondrial ROS and protects from toxicity of complex I inhibitors. (A) Measurement of complex I activity in control and AUL12-treated (2 μM) mitochondria. (B) Analysis of mitochondrial ROS levels by MitoSOX staining in control and AUL12-treated (4 μM, 1 hour) cells. (C) Cytofluorimetric assessment of cell viability following exposure to AUL12 (2/4 μM, 24h) with or without the antioxidant N-acetyl cysteine (NAC, 500 µM). (D) Analysis of mitochondrial ROS levels by MitoSOX staining in control and rotenone-treated (200 nM, 1 hour) cells. (E) Cell viability following exposure to rotenone (200/500 nM) with or without NAC (500 µM) was assessed as in (C). Data are reported as mean ± SD values (n ≥ 3); ***: p<0.001; **: p<0.01 and *: p<0.05 with a Student’s *t* test analysis. All experiments in the Figure were carried out on Nf1^+/+^ and Nf1^-/-^ MEFs. See also Figure S2.

### The alternative NADH dehydrogenase NDI1 increases oxygen consumption rate and confers resistance to oxidative stress in neurofibromin-expressing cells

In order to explore mechanistic links between complex I inhibition and downstream effects on cell viability and tumorigenicity, we bypassed complex I function by expressing NADH dehydrogenase 1 (NDI1), a single subunit enzyme from *S. cerevisiae* that catalyzes electron transfer from NADH to ubiquinone, without proton translocation and in a rotenone-insensitive way (Cannino et al., 2012; Seo et al., 1999). Expectedly, NDI1 made MEFs refractory to rotenone-dependent changes in ROS levels and oxygen consumption rate (OCR) (Figures S3A-S3C). OCR was partially inhibited in cells lacking neurofibromin downstream to Ras hyperactivation (Figure 3A, S3C and S3D). After NDI1 expression, Nf1^-/-^ MEFs increased their OCR and reached the level of Nf1^+/+^ cells, which instead were not affected by NDI1 (Figure 3A). Conversely, OCR was lowered by the complex I inhibitor AUL12 only in wild-type cells, making them reach the same level of Nf1^-/-^ MEFs, and NDI1 could circumvent this OCR inhibition (Figures 3B and 3C and Figures S3E and S3F). Together, these observations indicate that complex I inhibition is responsible for the lower respiratory rate of cells lacking neurofibromin with respect to Nf1^+/+^ cells. The presence of NDI1 also blunted the increase in mitochondrial ROS and the consequent toxicity of the complex I targeting compounds AUL12, EM20-25 and rotenone in Nf1^+/+^ MEFs (Figures 3D and 3E and Figures S3B, S3G and S3H; compare with Figures 2B-2E and Figure S2A).

**Figure 3.**
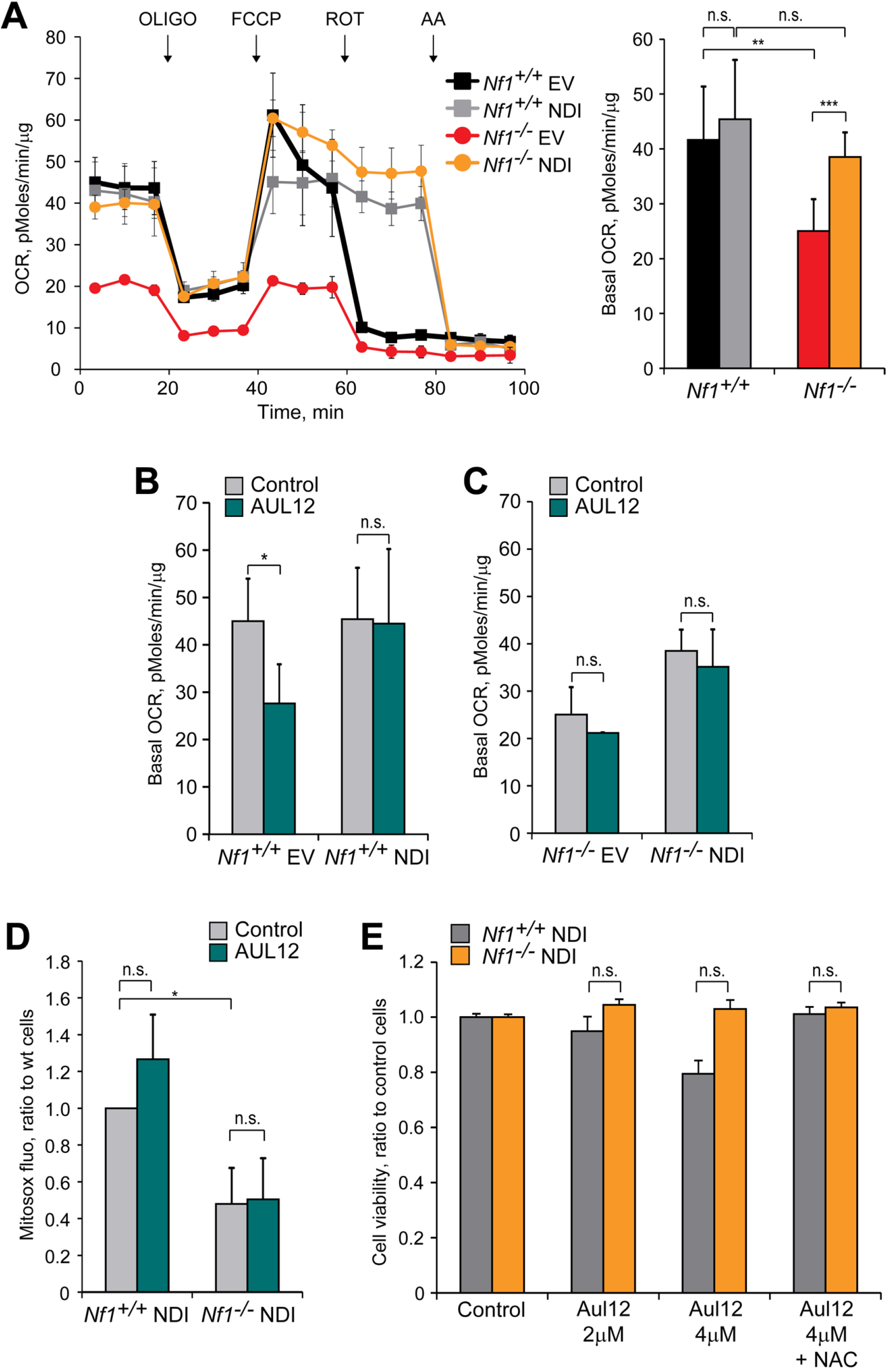
Effects of the alternative NADH dehydrogenase NDI1 on oxygen consumption rate (OCR), ROS and viability of cells. (A-C) Representative OCR traces (A) and quantification of basal OCR (A-C) of Nf1^+/+^ and Nf1^-/-^ MEFs. In (A), the ATP synthase inhibitor oligomycin (0.8 µM), the proton uncoupler carbonyl cyanide-4-(trifluoromethoxy)phenylhydrazone (FCCP, 1 µM) and the respiratory complex I and III inhibitors rotenone (0.5 µM) and antimycin A (1 µM), respectively, were added as indicated. In (B and C) AUL12 (4 µM) was added 45 minutes before recordings. NDI: cells expressing the pWPI-NDI1 construct; EV: cells expressing the pWPI empty vector. (D) Analysis of mitochondrial ROS levels by MitoSOX staining in control and AUL12-treated (4 μM, 1 hour) Nf1^+/+^ and Nf1^-/-^ MEFs expressing NDI1. (E) Cell viability assessment following exposure of Nf1^+/+^ and Nf1^-/-^ MEFs expressing NDI1 to AUL12 (2-4 µM, 24h) with or without NAC (500 μM). Data are reported as mean ± SD values (n ≥ 3); ***: p<0.001; **: p<0.01 and *: p<0.05 with a Student’s *t* test analysis. See also Figure S3.

### NDI1 impairs tumorigenicity of neurofibromin-deficient cells through SIRT3 reactivation

These data demonstrate that neurofibromin loss inhibits complex I activity via Ras-ERK signaling, leading to OCR down-regulation, lower mitochondrial ROS levels and resistance to oxidants targeting complex I. NDI1 expression can reverse this chain of events and constitutes a tool to investigate the impact of complex I inhibition on the tumorigenic potential of cells lacking neurofibromin.

NDI1 expression raised the intracellular NAD^+^/NADH ratio and NAD^+^ levels (Figure 4A and Figure S4A), in accord with its NADH dehydrogenase activity, and decreased tumorigenicity of Nf1^-/-^ cells (Figure 4B). Hence, we explored the possibility of a direct connection between increased NAD^+^/NADH balance and neoplastic growth driven by neurofibromin deficiency. The NAD^+^ precursors nicotinic acid (NIC) and nicotinamide (NAM), which augment NAD^+^ levels through the Preiss-Handler and the salvage pathway, respectively (Katsyuba and Auwerx, 2017), decreased tumorigenicity of Nf1^-/-^ cells to levels comparable to their NDI1-expressing counterparts, on which NIC and NAM were ineffective (Figure 4C and Figure S4B).

**Figure 4.**
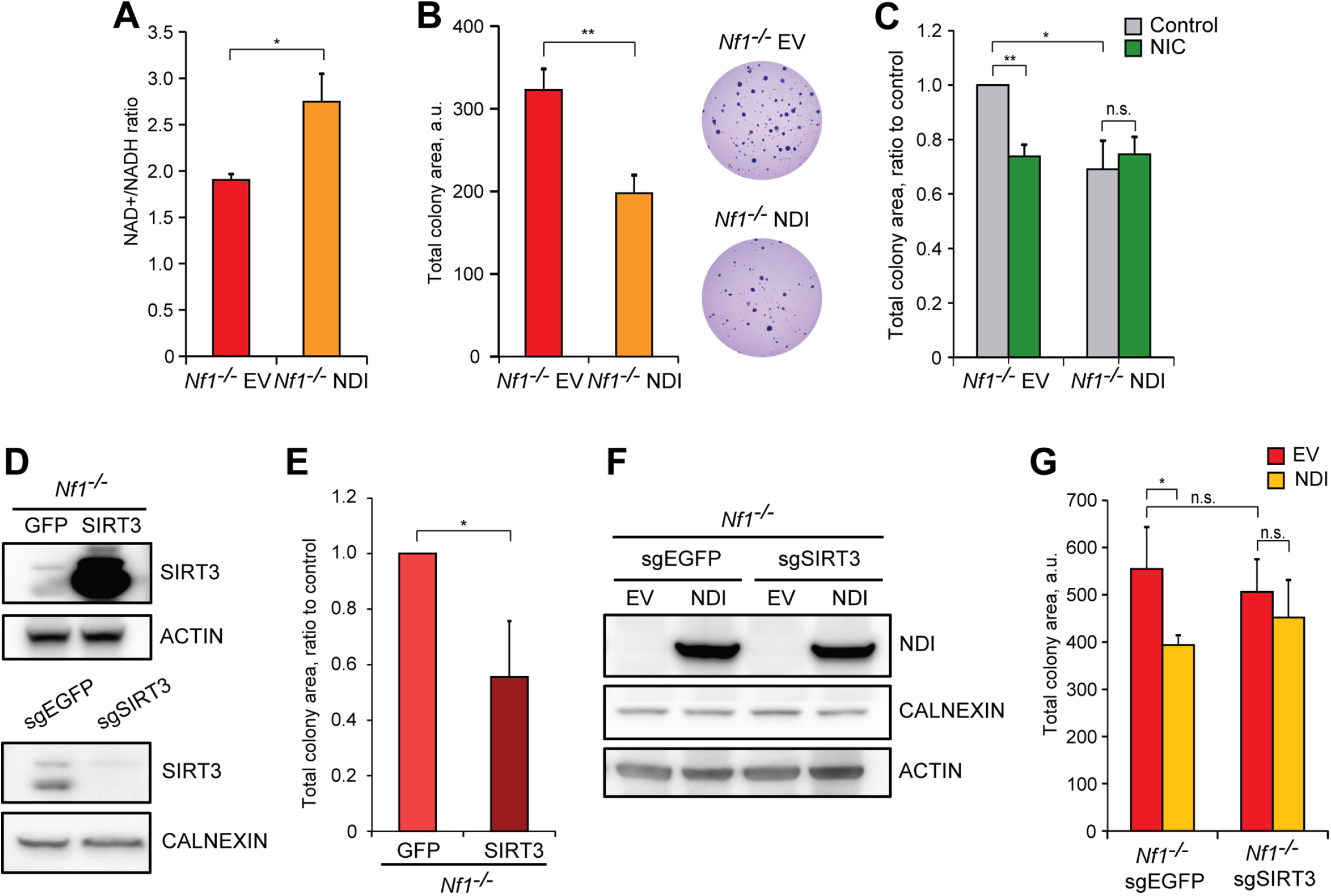
NDI1, nicotinic acid (NIC) and SIRT3 impair tumorigenicity of neurofibromin-deficient cells. (A) Spectrophotometric analysis of the NAD^+^/NADH ratio following NDI expression. (B) Measurement of colony area in soft agar experiments (left) and representative pictures of colonies (right) in Nf1^-/-^ cells expressing NDI. (C) Effect of NIC treatment (5 mM, 3 weeks; C) on colony growth in soft agar of Nf1^-/-^ cells. (D) WB analysis of SIRT3 protein levels after expression of the pFUGW-GFP/SIRT3 construct (upper part) or SIRT3 knocking out (sgSIRT3, lower part). Negative controls were cells expressing pFUGW-GFP (GFP, upper part) or scrambled single guide targeting EGFP (sgEGFP, lower part). (E) Effect of SIRT3 upregulation on soft agar colony formation of Nf1^-/-^ cells. (F) WB analysis of NDI1 expression in SIRT3 wild-type and knock-out (sgEGFP and sgSIRT3, respectively) neurofibromin-deficient cells. (G) Effect of NDI1 expression on soft agar growth of SIRT3 wild type (sgEGFP) and KO (sgSIRT3) Nf1^-/-^ MEFs. All experiments were carried out on Nf1^-/-^ MEFs. NDI: cells expressing the pWPI-NDI1 construct; EV: cells expressing the pWPI empty vector. In (D and F), actin and calnexin were used as loading controls. Data are reported as mean ± SD values (n ≥ 3); ***: p<0.001; **: p<0.01 and *: p<0.05 with a Student’s *t* test analysis. See also Figure S4.

Increased mitochondrial NAD^+^ levels can enhance the activity of the mitochondrial sirtuins SIRT3, SIRT4 and SIRT5. We found that SIRT3 overexpression in cells without neurofibromin (Figure 4D) decreased their tumorigenicity (Figure 4E), while SIRT4 or SIRT5 overexpression could not affect it (Figure S4C). Knocking-out SIRT3 (Figure 4D) did not change the ability of Nf1^-/-^ cells to form colonies but made them insensitive to the anti-neoplastic effect of NDI1 (Figures 4F and 4G). The key antioxidant enzyme SOD2 is a *bona fide* indicator of SIRT3 activity, as it is activated by SIRT3-dependent deacetylation (George and Ahmad, 2016; Tao et al., 2010). We found a decreased SOD2 acetylation after NDI1 expression (Figure S4D), where the protein levels of SOD2 and of other SIRT3 targets were unchanged (Figures S4D and S4E). In accord with the hypothesis of a pathway connecting increased intracellular NAD^+^/NADH ratio with anti-neoplastic SIRT3 induction, NIC/NAM administration or SIRT3 overexpression were able to reduce SOD2 acetylation status (Figures S4F and S4G), and forcing SOD2 activity through its overexpression decreased the tumorigenic potential of neurofibromin-deficient cells (Figure S4H). To further corroborate this model we analyzed cell types derived from MPNSTs, malignancies typically associated to NF1 that are endowed with a profound metabolic rewiring (Lin and Gutmann, 2013) and for which effective therapies are lacking (Reilly et al., 2017). NDI1 expression increased OCR (Figure 5A and Figures S5A and S5B) and reduced tumorigenicity (Figure 5B) in MPNST cell models, as well as SIRT3 overexpression (Figures 5C and 5D).

**Figure 5.**
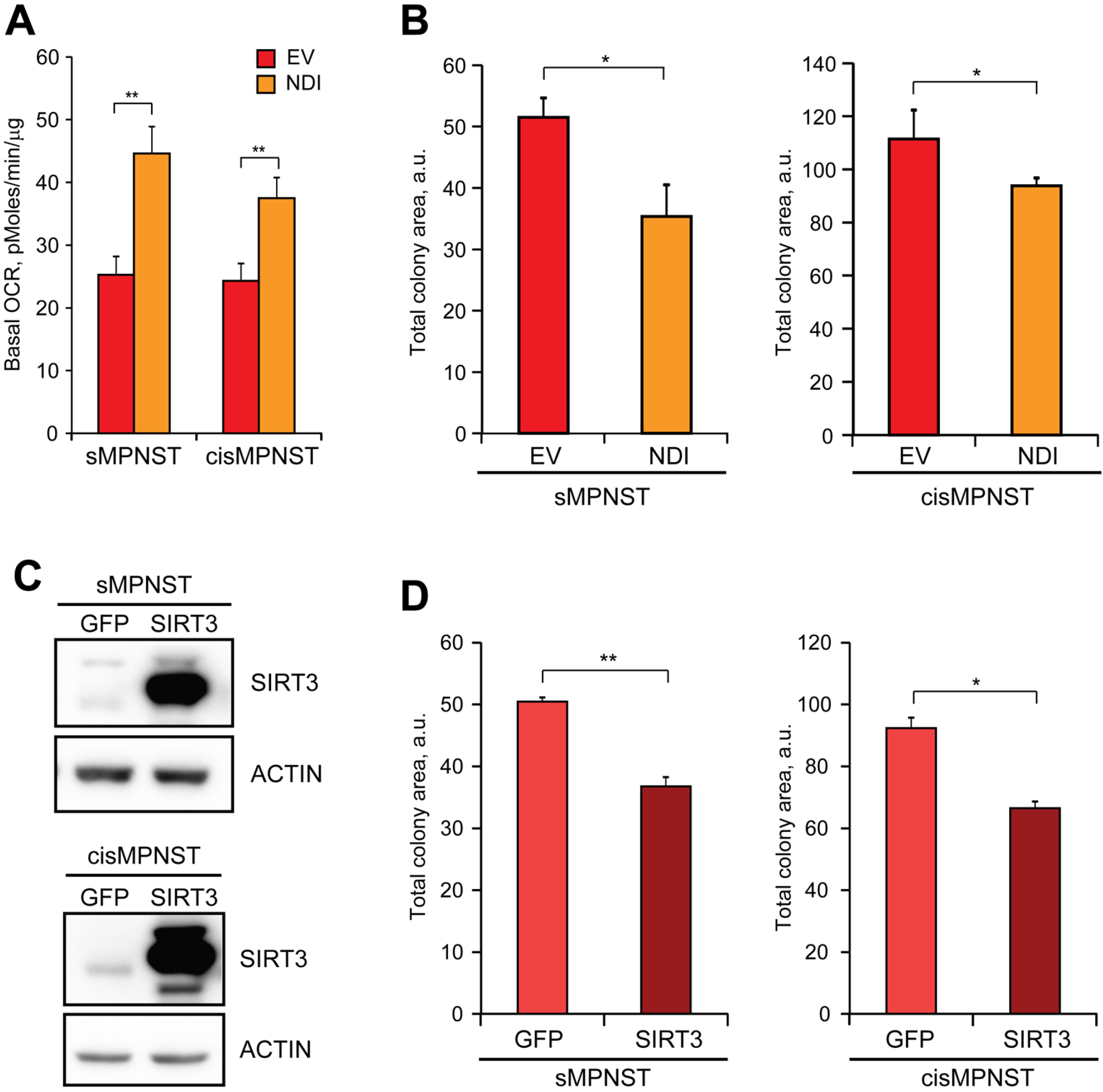
Effect of NDI or SIRT3 expression on MPNST cell tumorigenicity. (A and B) Effect of NDI1 expression on OCR (A) and soft agar colony formation (B) of MPNST cells. (C and D) SIRT3 over-expression after transfection with pFUGW-SIRT3 (C) decreased Matrigel colony formation of MPNST cells (D). In (C), actin was used as a loading control. All along the Figure, the MPNST cell models sMPNST and cisMPNST were used. NDI1: cells expressing the pWPI-NDI1 construct; EV: cells expressing the pWPI empty vector; SIRT3: cells expressing pFUGW-SIRT3; GFP: cells expressing pFUGW-GFP. Data are reported as mean ± SD values (n ≥ 3); ***: p<0.001; **: p<0.01 and *: p<0.05 with a Student’s *t* test analysis. See also Figure S5.

These results indicate that, upon neurofibromin loss, hyperactivation of Ras/MEK/ERK signaling inhibits complex I, causing a decrease in the NAD^+^/NADH ratio and the ensuing SIRT3 repression. Raising the NAD^+^/NADH ratio *via* NDI1 expression has an anti-neoplastic effect by enhancing SIRT3 activity.

### TRAP1 ablation and increased NAD^+^ levels/SIRT3 activity exert anti-neoplastic bioenergetic effects on MPNST growth

We have previously shown that the mitochondrial chaperone TRAP1 has a pro-neoplastic effect by inhibiting SDH activity (Sciacovelli et al., 2013) and this also applies to NF1-related models, where TRAP1 activity is enhanced in an ERK-dependent way (Masgras et al., 2017a). SDH constitutes a potential point of intersection between the bioenergetic effects caused by SIRT3 induction and TRAP1 ablation as SIRT3 increases SDH activity (Cimen et al., 2010; Finley et al., 2011b) similarly to TRAP1 absence or inhibition (Masgras et al., 2017a; Sanchez-Martin et al., 2020a). However, it was also proposed that TRAP1 is activated by SIRT3-dependent deacetylation in a glioblastoma model (Park et al., 2019), making it difficult to draw a comprehensive picture.

We therefore explored whether there is any interplay between the pro-tumoral roles exerted by TRAP1 and by SIRT3 inhibition downstream to complex I inhibition in NF1-related malignant cells. We found that neither TRAP1 knock-out nor SIRT3 overexpression change protein levels of SDH subunits (Figure 6A), but both conditions raised SDH activity to a similar extent and without any additive effect (Figure 6B), which is strongly suggestive of a common effector mechanism. Succinate oxidation was similarly enhanced in MPNST cells upon NIC administration in *in vitro* conditions of neoplastic growth (Figure 6C) without any change in SDH protein levels (Figure 6D), and NIC increased the anti-tumor effect of knocking-out TRAP1 expression (Figure 6E). In further accord with a synergistic antineoplastic effect of boosting NAD^+^ levels and ablating TRAP1 expression, administration of QS10, a short chain quinone that acts as an electron acceptor for NADH, reduced colony formation with a higher efficacy in TRAP1 knock-out than in TRAP1-expressing sMPNST cells (Figure S6A).

**Figure 6.**
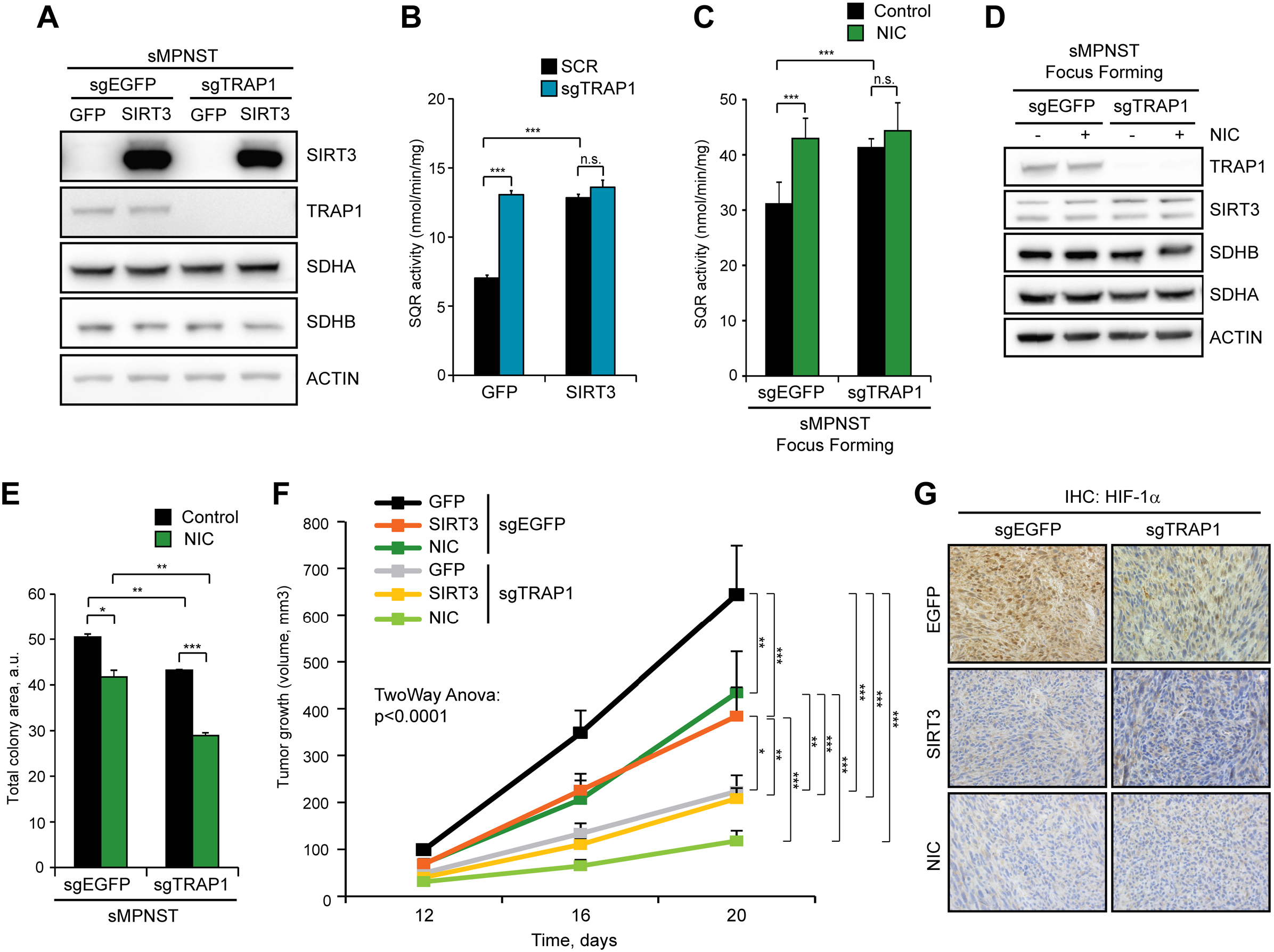
Anti-neoplastic effects of SIRT3 and NIC and of TRAP1 ablation. (A) WB analysis of SIRT3 and SDHA/B protein levels. Actin was used as a loading control. (B) Analysis of succinate-coenzyme Q reductase (SQR) activity of SDH in control and TRAP1 knock-out (sgTRAP1) sMPNST cells. SIRT3: cells expressing pFUGW-SIRT3; GFP: cells expressing pFUGW-GFP. (C-E) Effect of NIC treatment (5 mM) on SQR activity (C), SDHA/B protein levels (D) or Matrigel colony formation (E) of control and TRAP1 knock out sMPNST cells. Measurements in (C, D) were carried out on cells undergoing tumorigenic growth (day 5^th^ of focus forming assay). In (D) actin was used as a loading control. (F) Kinetics of tumor growth of control and TRAP1 knock-out sMPNST cells xenografted in nude mice. Cells are labeled as in (A). Where indicated, animals were supplied with NIC (1 % in drinking water). Tumor volume is expressed in mm^3^. (G) Immunohistochemical analysis of HIF1α expression in tumor samples from (E). 40X magnification. In (B, C, E) data are reported as mean ± SD values (n ≥ 3) and analyzed using a Student’s *t* test. In (F) data are reported as mean ± SEM values (n ≥ 7) and analyzed by a two-way ANOVA followed by Bonferroni post-test. ***: p<0.001; **: p<0.01 and *: p<0.05. See also Figure S6.

*In vivo*, neoplastic growth of MPNST cells was inhibited by SIRT3 overexpression or treatment of mice with NIC, as well as by TRAP1 ablation (Figure 6F and Figure S6B). SIRT3 overexpression did not further decrease tumor growth in a TRAP1-null background, whereas NIC treatment almost ablated it (Figure 6F and Figure S6B). The stabilization of the transcription factor HIF1α is crucial to the pro-neoplastic function of TRAP1 (Sciacovelli et al., 2013). We found that TRAP1 ablation, SIRT3 induction and NIC treatment markedly reduced the nuclear localization of HIF1α in xenografted sMPNST cells (Figure 6G). In tumor samples both the NAD^+^/NADH ratio and NAD^+^ were increased following NIC administration and when TRAP1 was ablated, and were dramatically raised when NIC was provided to animals harboring TRAP1 knock-out cells (Figure S6C-D). These observations are in accord with a global induction of the OXPHOS activity of the cells caused by TRAP1 inhibition (Rasola et al., 2014), suggesting that its anti-tumor effect could depend not only on induction of SDH, but also of complex I. Moreover, a high NAD^+^/NADH ratio might have multiple effects on the biology of the cell (Katsyuba et al., 2020), explaining the enhanced anti-neoplastic efficacy of NIC administration on TRAP1 knock-out cells.

## DISCUSSION

In the present study, we demonstrate that deficiency of the RasGAP neurofibromin down-modulates respiration. This specifically occurs through inhibition of NADH dehydrogenase following hyper-activation of Ras-MEK-ERK pathway that is mandatory for growth of NF1-related tumors (Ratner and Miller, 2015). As a result, cells change redox homeostasis and unbalance their NAD^+^/NADH ratio with pro-neoplastic consequences.

Previous reports of respiratory complex I inhibition in K-Ras transformed cells (Baracca et al., 2010; Hu et al., 2012) have suggested that it could contribute to cancer-associated biological routines such as proliferation, resistance to cell death and metastatic growth (Urra et al., 2017). Consistently, several molecules inhibiting complex I have been proposed as potential anticancer agents (Vatrinet et al., 2015). Here we detail the oncogenic function of complex I inhibition, showing that it has a dual effect that can be exploited by the tumor cell. On the one hand, it keeps intracellular ROS levels at bay, shielding cells from a variety of environmental insults that can hamper neoplastic growth by eliciting oxidative stress. These include several chemotherapeutics, but also hypoxic conditions frequently encountered by cells during the turbulent growth of the tumor mass (Sabharwal and Schumacker, 2014). It follows that anti-neoplastic strategies alternative to treatments based on pro-oxidants must be adopted for circumventing this phenotype. On the other hand, a low NADH dehydrogenase activity means dropping the NAD^+^/NADH ratio, with widespread consequences on the metabolic equilibrium of the cell. NAD^+^ acts as an electron acceptor in a variety of biochemical reactions that encompass glycolysis, oxidative decarboxylation of pyruvate to acetyl-CoA, β-oxidation of fatty acids and TCA cycle (Katsyuba et al., 2020). In addition, NAD^+^ can regulate metabolism by acting as a co-substrate for sirtuin deacylases (Chalkiadaki and Guarente, 2015). We find that increasing the NAD^+^/NADH balance or overexpressing the NAD^+^-dependent mitochondrial deacetylase SIRT3, but not SIRT4 or SIRT5, hinder tumorigenicity in neurofibromin-deficient cells, including highly aggressive MPNST models. Conversely, knocking-out SIRT3 does not further increase MPNST tumorigenicity, indicating that it is constitutively inhibited in neurofibromin-lacking cells. These observations point toward a *bona fide* tumor suppressor role for SIRT3, which is abrogated by respiratory complex I inhibition and the ensuing decrease of NAD^+^/NADH ratio in NF1-related tumor cells.

The use of the yeast NADH dehydrogenase NDI1, a rotenone insensitive, single subunit protein capable of restoring NADH oxidation and mitochondrial respiration in cells devoid of complex I activity (Cannino et al., 2012), increases respiration of Nf1^-/-^ cells to the levels of their neurofibromin-expressing counterparts. It was reported that NDI1 impairs tumorigenicity of cells carrying a complex I dysfunction (Sharma et al., 2011) and in a breast cancer model (Santidrian et al., 2013). Accordingly, we observe that NDI1 thwarts tumorigenicity of NF1-related tumor cells, directly demonstrating a causal connection between a decrement in the activity of respiratory complex I and neoplastic growth. NDI1 was ineffective on the tumorigenic potential of cells where we knocked-out SIRT3 expression, further underlining that NDI1 hampers neoplastic growth in a SIRT3-dependent manner.

SIRT3 works as a master regulator of mitochondrial metabolism and redox homeostasis by affecting the activity of enzymes such as glutamate dehydrogenase, isocitrate dehydrogenase 2, serine hydroxymethyltransferase 2, superoxide dismutase and pyruvate dehydrogenase (Chalkiadaki and Guarente, 2015; van de Ven et al., 2017). Moreover, it has been proposed that SIRT3 activates the mitochondrial chaperone TRAP1, thus contributing to the maintenance of cancer stem cells in a glioblastoma model (Park et al., 2019). We have formerly demonstrated that TRAP1 is pro-neoplastic in diverse neoplastic models by inhibiting SDH activity, thus activating HIF1α (Guzzo et al., 2014; Kowalik et al., 2016; Masgras et al., 2017b; Rasola et al., 2014; Sciacovelli et al., 2013), and that a mitochondrial fraction of ERK (Rasola et al., 2010) phosphorylates TRAP1 and enhances its tumorigenic activity in NF1 models (Masgras et al., 2017a). Consequently, it is difficult to reconcile an oncogenic function of TRAP1 with its activation by SIRT3, if the latter plays a tumor suppressor role. In this regard, we have recently found that SIRT3 overexpression raises SDH activity, mimicking the absence of TRAP1 (Sanchez-Martin et al., 2020a). In this study, we report that knocking-out TRAP1 expression is ineffective in further increasing SDH activity of SIRT3-overexpressing cells, in accord with SIRT3 enhancing SDH activity via TRAP1 inhibition. Therefore, we propose that SIRT3 activation in MPNST cells is anti-neoplastic at least in part by re-establishing SDH activity and counteracting HIF1α stabilization, as we have observed in xenografted cancer cells. Given that both SIRT3 overexpression and its induction through agents increasing NAD^+^ (NIC/NAM administration) activate SOD2, and that SOD2-overexpressing cells decrease their tumorigenicity, it is possible that SIRT3-mediated activation of SOD2 further contributes in the repression of neoplastic growth of Nf1^-/-^ cells.

We envision that enhancement of SIRT3 activity and replenishment of NAD^+^ levels result in a multifaceted metabolic rewiring, which can oppose NF1-related cancer growth by affecting multiple bioenergetic pathways. Accordingly, the anti-neoplastic effect of both SIRT3 overexpression and NIC administration is higher in TRAP1 knock-out cells, suggesting that targeting multiple metabolic components can dramatically hit tumor growth. Indeed, NAD^+^ levels increased the most following NIC treatment in a TRAP1-null background. These data open obvious therapeutic perspectives, and the recently identified selective inhibitors of TRAP1 (Sanchez-Martin et al., 2020a; Sanchez-Martin et al., 2020b; Sanchez Martin et al., 2020c) are interesting candidates as anti-neoplastic leads in NF1-related tumors.

Key findings of the present work are the identification of a complex bioenergetic adaptation in mitochondria of neurofibromin-deficient cells and the demonstration that this rewiring in respiratory function contributes to the tumorigenic potential of NF1-related neoplasms. These observations constitute a conceptual starting point for drawing antineoplastic strategies based on combinatorial targeting multiple metabolic hubs in tumor models endowed with oncogenic Ras/ERK signaling.

### Limitations

This study discloses the contribution of a decreased respiration in the tumorigenic process of cells lacking neurofibromin, observing that a concomitant inhibition in the activity of respiratory complex I and of succinate dehydrogenase (SDH, respiratory complex II) connects to tumor growth through the unbalance of the NAD^+^/NADH ratio and the consequent inhibition of the SIRT3 deacetylase. This impinges on SDH inhibition mediated by the mitochondrial chaperone TRAP1, leading to the ensuing pro-neoplastic stabilization of HIF1α. However, we cannot exclude that changes in other biochemical pathways can irradiate from NAD^+^/NADH unbalance and SIRT3 inhibition, further supporting the tumorigenic process in neurofibromin-deficient cells. We also observe that inhibition of respiratory complex I decreases mitochondrial ROS levels in NF1-related cells, making them more resistant to toxicity mediated by several pro-oxidants. Yet, the interplay between redox balance and neoplastic progression is complex and context-dependent, suggesting that a low level of ROS generation at complex I could have multifaceted roles in different stages of neurofibroma growth, which should be thoroughly investigated *in vivo*. It also remains to be elucidated whether our findings of a respiratory inhibition downstream to the Ras/MEK/ERK signaling pathway are found in other tumor models characterized by hyperactivation of this pathway, and how they contribute to the neoplastic process in these settings.

## Supporting information

Supplemental information contains 6 supplemental figures and 1 supplemental table

## ACKNOWLEDGMENTS

A.R. was supported by grants from University of Padova, Neurofibromatosis Therapeutic Acceleration Program and Associazione Italiana Ricerca Cancro (AIRC grant IG 2017/20749). I.M. and F.C. are recipients of Young Investigator Award Grants from Children’s Tumor Foundation. We thank Reuven Stein for MEFs; Eric Dufour for pWPI-NDI1 vectors; Danica Chen for pFUGW-SIRT3 and pFUGW-SOD2 vectors; Dolores Fregona for AUL12 compound; Elena Trevisan for excellent technical assistance, and Paolo Bernardi for helpful comments and discussion on the project.

## AUTHOR CONTRIBUTIONS

Conceptualization: I.M. and A.R.; visualization, I.M.; methodology: I.M., G.C. and A.C.; investigation: I.M., G.C., F.C., C.S.M., M.P., A.M.; formal analysis: I.M., G.C., F.C., C.S.M., M.P. and A.M.; resources: G.C., M.P.; writing-original draft: I.M. and A.R.; writing-review and editing: I.M., A.C. and A.R.; funding acquisition; A.R.; project administration: A.R.; supervision: A.R.

## MATERIALS AND METHODS

### LEAD CONTACT

Further information and requests for resources and reagents should be directed to and will be fulfilled by the Lead Contact Andrea Rasola.

## EXPERIMENTAL MODEL AND SUBJECT DETAILS

### Nf1^-/-^ MEFs and MPNST cell lines

Mouse embryonic fibroblasts (MEFs) were derived from both wild-type mice and syngenic Nf1-knockout animals (Nf1^+/+^ and Nf1^-/-^ MEFs, respectively) (Shapira et al., 2007) and were kindly provided by Dr. R. Stein, University of Tel Aviv, Ramat Aviv, Israel. sMPNST cells were established from neurofibromin 1 (Nf1) and p53-deficient skin precursors (SKP) (Mo et al., 2013); cisMPNST cells were derived from spontaneous MPNSTs arising in cis Nf1^+/-^;P53^+/-^ mice (Vogel et al., 1999); both mouse MPNST cells lines were kindly provided by Dr. Lu Q. Le, University of Texas Southwestern Medical Center, Dallas, TX. Human plexiform neurofibroma ipNF 04.4 cells were generated and provided by Dr. Margaret R. Wallace, University of Florida, College of Medicine, Gainesville, FL (Li et al., 2016). All cells were grown in Dulbecco’s modified Eagle’s medium (DMEM) supplemented with 10% fetal bovine serum, 2 mM glutamine, 1 mM sodium pyruvate and 100 µg/ml penicillin and streptomycin at 37°C in a humidified atmosphere containing 5% CO_2_.

### Mice

We employed nude female mice (Charles River). All mice were housed on a 12:12 hours light:dark cycle at 25°C. Six-week-old animals (n = 7) were subjected to tumor grafts by subcutaneous injection of sMPNST cells. All animal procedures were in accordance the animal care committee at University of Padova and the Italian Ministry of Health.

### Generation of TRAP1 and SIRT3 knock-out cell lines

TRAP1 knock-out cells were generated by using the clustered regulatory interspaced short palindromic repeat (CRISPR)-Cas9 gene system (Sanjana et al., 2014). Sequences for the single guides (for mouse TRAP1: 5’-CACCGCGCCGAACTCCAGCCAGCGC-3’ and 5’-CACCGTTTGTGTGGGGCCCCTAAAC-3’; for mouse SIRT3: 5’-CACCGTCTATACACAGAACATCGAC-3’ and 5’-CACCGTTGCTGTAGAGGCCGCTCCC-3’ and 5’-CACCGACATTGGGCCTGTAGTGCCC-3’) were obtained by using the CRISPR design tool (http://www.crispr.mit.edu). Scrambled single guide targeting EGFP gene were used as negative controls. Oligonucleotide pairs were annealed and cloned into the transfer plasmid lentiCRISPRv2 (Addgene, #52961) and co-transfected with the packaging plasmids pMDLg/pRRE (Addgene, #12251), pRSV-Rev (Addgene, #12253) and pMD2.G (Addgene, #12259) into human embryonic kidney (HEK) 293T cells for viral production. Recombinant virus was collected and used to infect cells by standard methods. Infected cells were then selected with 2 µg/ml puromycin.

### Generation of overexpressing cell lines

NF1-GRD sequence was introduced in Nf1^-/-^ MEFs using the pMSCV-GRD vector (Shapira et al., 2007). pBABE vectors were used for expression of constitutively active (CA: S217E/S221E) and dominant negative (DN: S217A) MEK1 and hyperactive RAS (G12D mutant) (Addgene, #58902). For SIRT3, SIRT4 and SIRT5 overexpression, pcDNA3.1-SIRT3/SIRT4/SIRT5 (Addgene, #13814/13815/13816) were used for sub-cloning sirtuin genes into a pBABE vector. pWPI vector was used for NDI1 expression (Cannino et al., 2012) and pFUGW was used for overexpression of GFP (as a control) SIRT3 and SOD2 (Qiu et al., 2010). Retroviral or lentiviral vectors were used for the production of viral particles and infected cells were then selected with 2 µg/ml puromycin.

### Mitochondria isolation

Mitochondria were isolated after cell disruption with a glass-Teflon or electrical potter (Sigma) in a buffer composed of 250 mM sucrose, 10 mM Tris-HCl, 0.1 mM EGTA-Tris, pH 7.4. Nuclei and plasma membrane fractions were separated by a first mild centrifugation (700 g, 10 min); mitochondria were then spinned down at 7000 g, 10 min, and washed twice (7000 g, 10 min each). All procedures were carried out at 4°C.

### Western Immunoblots and Immunoprecipitations

For Western immunoblots analyses, cells or isolated mitochondria were lysed at 4°C in a buffer composed of 150 mM NaCl, 20 mM Tris-HCl pH 7.4, 5 mM EDTA, 10% glycerol, 1% Triton X-100 (lysis buffer), in the presence of phosphatase and protease inhibitors (Sigma). Lysates were then cleared with a centrifugation at 18000 *g* for 30 minutes at 4°C, and proteins were quantified using a BCA Protein Assay Kit (Thermo Scientific-Pierce).

Protein immunoprecipitations were carried out on 200 µg isolated mitochondria. Lysates were pre-cleared with an incubation with Dynabeads® Protein G (Thermo Fisher Scientific) for 1 hour at 4°C and then incubated in agitation for 18 h at 4°C with the antibody conjugated to fresh Dynabeads® Protein G. Where indicated, an unrelated anti mouse IgG was added as a negative isotype control. Beads were then washed several times in lysis buffer.

Proteins extracted from total cell or mitochondrial lysates or from immunoprecipitations were then boiled for 5 minutes in Laemmli sample buffer, separated in reducing conditions on SDS-polyacrylamide gels and transferred onto Hybond-C Extra membranes (Amersham) following standard methods. Primary antibodies were incubated 16 hours at 4°C, and horseradish peroxidase-conjugated secondary antibodies were added for 1 hour at room temperature. Proteins were visualized by enhanced chemiluminescence (Millipore).

### Blue native polyacrylamide gel electrophoreses (BN-PAGE)

BN-PAGE experiments were performed on mitochondria isolated as described above. ETC complexes and super-complexes were extracted at 4°C for 2 minutes in the presence of 2% *n*-dodecyl-β-D-maltoside (DDM) and 2% digitonin, respectively, starting from 200 μg of mitochondria in a buffer composed of 1 M aminocaproic acid, 50 mM Bis Tris pH 7. After extraction mitochondria were spinned at 100000 *g* for 30 minutes and supernatants were collected and loaded on polyacrylamide Native-PAGE 3-12% Bis-Tris gradient gels (Invitrogen) after addition of Coomassie Blue G250 (Invitrogen). Protein complexes were then visualized after 18 hours of Coomassie Blue G-250 staining and/or subjected to in gel activity assay (see below). Bands corresponding to the indicated respiratory chain complexes were cut and subjected to 2D-SDS-PAGE, in order to separate single protein components, which were identified by Western immunoblotting.

### Measurements of NADH dehydrogenase activity

Mitochondrial enriched fractions (20-40 µg per trace) were used for spectrophotometric recordings of NADH dehydrogenase activity of mitochondrial complex I. The rotenone-sensitive NADH-CoQ oxidoreductase activity was detected following the decrease in absorbance due to the oxidation of NADH at 340 nm (ε=6.2 mM^-1^ cm^-1^). Reaction was performed at 30°C in 10 mM Tris-HCl pH 8 buffer containing 5 µM alamethicin, 3 mg/ml BSA, 5 µM sodium azide, 2 µM antimycin A, 65 µM coenzyme Q1, and 100 µM NADH. The NADH-ubiquinone oxidoreductase activity was measured for 3-5 min before the addition of rotenone (10 µM), after which the activity was measured for an additional 3-5 min. Measurements of complex I activity were normalized for citrate synthase (CS) activity. To measure CS activity, citrate formation was determined with a spectrophotometer as an increase in absorbance at 420 nm at 37°C (ε=13.6 mM^-1^ cm^-1^). Reaction buffer was composed of 100 mM Tris-HCl pH 8, 0.1% Triton X-100, 100 μM 5,5’-dithiobis-(2-nitrobenzoic acid) (DTNB), 300 μM acetyl -CoA, and 500 μM oxaloacetate. In gel complex I activity was performed by incubating Blue Native gels overnight at room temperature with a solution composed of 2 mM Tris–HCl, pH 7.4, 0.1 mg/ml NADH, and 2.5 mg/ml NTB (nitrotetrazolium blue).

### SDH succinate:coenzyme Q reductase (SQR) activity

To measure the SQR enzymatic activity of succinate dehydrogenase (SDH), cells were collected at 4°C in a buffer composed of 25 mM potassium phosphate, pH 7.2, 5 mM magnesium chloride and protease and phosphatase inhibitors. Cell homogenates (40 µg protein per trace) were then pre-incubated for 10 minutes at 30°C in a buffer containing 25 mM potassium phosphate, pH 7.2, 5 mM magnesium chloride, 20 mM sodium succinate and 10 µM alamethicin. After the pre-incubation time, 2 µM rotenone, 5 µM antimycin A and 5 mM sodium azide were added to the medium. Reaction was started after the addition of 100 µM 2,6-dichloroindophenol (DCPIP) and 65 µM coenzyme Q1. SQR enzymatic activity was recorded following the reduction of DCPIP at 600 nM (ε = 19.1 mM-1 cm-1) for 20 min at 30°C. Each measurement of SDH activity was normalized for protein amount.

### Oxygen Consumption Rate (OCR) experiments

OCR was assessed in real-time with the XF24 Extracellular Flux Analyzer (Seahorse Biosciences). Cells (2×10^4^/well) were plated the day before the experiment in a DMEM/10% serum medium; experiments were carried out on confluent monolayers. Before starting measurements, cells were placed in a running DMEM medium (supplemented with 25 mM glucose, 2 mM glutamine, 1 mM sodium pyruvate, and without serum and sodium bicarbonate) and pre-incubated for 1 hour at 37°C in atmospheric CO_2_. OCR values were then normalized for the protein content of each sample. An accurate titration with the uncoupler FCCP was performed for each cell type, in order to utilize the FCCP concentration (0.5-1 µM, depending on the cell type) that maximally increases OCR.

### Measurement of the NAD^+^/NADH ratio

To measure the NAD^+^/NADH ratio in mitochondria, cells were cultured in standard conditions prior to mitochondria isolation that was performed as described above. The NAD^+^/NADH ratio was measured using the NAD^+^/NADH assay kit (Colorimetric, Abcam), according to the manufacturer’s instructions. Briefly, isolated mitochondria were lysed by two freeze/thaw cycles in NAD^+^/NADH Extraction Buffer and vortexed for 10 seconds. After centrifugation, half supernatant was used for measurement of NADt (total amount of NAD^+^ and NADH), and the other half was used for measurement of NADH (after decomposition of NAD^+^ at 60°C for 30 minutes). Samples were incubated for 5 minutes with NAD Cycling Mix, followed by NADH Developer solution for 4 hours. Absorbance at 450 nm was then measured. The NAD^+^/NADH ratio was calculated as follows: (NADt -NADH)/NADH.

### Mass spectrometry analysis

NAD^+^/NADH ratio was calculated on tumor samples by mass spectrometry-based analysis. Tumor tissue specimens were flash frozen, weighted, and homogenized in 1 mL of 80% methanol. Samples were centrifuged at 20.000xg for 10 min at 4°C and the supernatants were transferred in a clean tube and dried using Speedvac. Protein pellet were kept for BCA/protein assay to be used for normalization. Dried samples were reconstituted with mQ water and frozen at -80°C. Mass spectrometry analysis was carried out with an LC-MS/MS (Quattro Premier interfaced with an Acquity UPLC system, Waters). The multiple reaction monitoring transition monitored for NAD^+^ was m/z 664.2 > 428.2 and for NADH m/z 666.2 > 649.1. Chromatographic resolution of NAD^+^ and NADH was achieved using an Atlantis dC18 column (2.1 150 mm, 5-m particle size, Waters) eluted with a linear gradient from 100% 10 mM ammonium formate (initial phase) to 10% 10 mM ammonium formate/90% methanol (Todisco et al., 2006). The flow was set at 0.3 ml/min. Calibration curves were established using standards, processed in the same conditions as the samples, at four concentrations (Frelin et al., 2012; Palmieri et al., 2014). The lines of best fit were determined using regression analysis based on the peak area of the analytes.

### Quantitative RT-PCR

Total RNA was isolated from cells using TRIzol reagent according to the manufacturer’s instructions. 2 µg of total RNA was used to synthesize cDNA with the SuperScript III First-Strand Synthesis System (Thermo Fisher Scientific) according to the manufacturer’s protocol. Quantitative RT-PCR was performed with the Biorad qRT-PCR machine using SYBR Green (Invitrogen). All reactions were performed for at least 6 biological replicates and the values expressed as fold increase in mRNA levels relative to control cells. Βeta-actin was used as a housekeeping gene. qRT-PCR primers are listed in Table S1.

### Measurement of ROS

Measurement of mitochondrial ROS were performed by MitoSOX (Thermo Fisher Scientific) staining according to manufacturer’s instructions followed by flow cytometry recordings. Briefly, cells were incubated with 2.5 μM MitoSOX for 15 minutes at 37°C in DMEM media depleted of FBS; next, treatments (e.g. rotenone, AUL12, antimycin A, etc.) were added and kept for the following 30-45 minutes. Then, cells were detached, resuspended in a buffer containing 135 mM NaCl, 10 mM HEPES and 5 mM CaCl_2_ (FACS mix solution) and analyzed. Changes in forward and side light scatter were assessed at the same time to measure alterations in cell dimension and granularity, respectively. Samples were analyzed on a FACSCanto II flow cytometer (Becton Dickinson). Data acquisition and analysis were performed using FACSDiva software.

### Cell viability assays

Cell viability was assessed either by a colorimetric MTS assay (Cell Titer 96® Aqueous One Solution; Promega) or by flow cytometry analysis. For MTS measurements, plates were incubated at 37°C overnight and read in a microplate spectrophotometer (Infinite® 200 PRO). Flow cytometry recordings were performed as described previously (Ciscato et al., 2014; Guzzo et al., 2014). Briefly, cells were stained with FITC-conjugated Annexin-V and 7-Aminoactinomycin D (7-AAD) to determine phosphatidylserine exposure on the cell surface (increased FITC-conjugated Annexin-V staining) and loss of plasma membrane integrity (7-AAD permeability and staining). Cells were incubated at 37°C in an assay buffer containing 135 mM sodium chloride, 10 mM HEPES, 5 mM calcium chloride and samples were then analyzed on a FACS Canto II flow cytometer (Becton Dickinson). Data acquisition and analysis were performed using FACSDiva software.

### *In vitro* tumorigenesis assays

Focus forming assays were performed on cells grown in 12-well culture plates in DMEM medium supplemented with 10% fetal bovine serum. When cells reached sub-confluence, serum concentration was reduced to 1% and NIC was added at a concentration of 5 mM. At the 3rd or 5th day after serum decrease, cells were scraped and collected at 4°C and SQR enzymatic activity of SDH was measured as described above. For the soft agar assay, cells were grown in 24 well plates covered by a bottom layer composed by DMEM medium mixed with low melting point agarose (Promega) at a final concentration of 1%, and by a top layer of DMEM medium supplemented with 1% serum and mixed with low melting point agarose at a final concentration of 0.6%. Cells (0.2×105/cm^2^) were added during the preparation of the upper layer, where they remained embedded. Dishes were then maintained in a humidified atmosphere of 5% CO_2_-95% air at 37°C for three weeks, adding medium (DMEM 2% serum) on the top of the two layers every 3^rd^ day. At the 25-30^th^ day, dishes were washed in PBS and colonies were stained with Crystal Violet 0.005% and analyzed with ImageJ software. Growth in 4% Matrigel (Corning) was performed in low adhesion 24 well plates in DMEM medium supplemented with 2% FBS. Cells (0.1×10^5^/cm^2^) were seeded in 600 μl final volume and after 2-3 days colonies were stained with Crystal Violet 0.005% and analyzed with ImageJ software.

### *In vivo* tumorigenesis assays

Experiments were performed in 8-week-old nude mice (Charles River, Wilmington, MA) treated in accordance with the European Community guidelines. Six mice were injected subcutaneously bilaterally in the flanks with 1.5×10^6^ sMPNST in 100 μl of serum-free sterile PBS mixed with 4% Matrigel. Nicotinic acid treatment (1% into drinking water) was administrated at day 2 following xenograft injection and refreshed every 3 days. Tumors were visible under the skin after 7-9 days and measured with caliper every four days (two major axes). Tumor volume was calculated using the formula: (a*b^2^)/2. After three weeks, mice were sacrificed and tumors stored at -80°C or fixed in formaldehyde and maintained in 70% ethanol for immunohistochemical analyses.

### Immunohistochemical analyses

Histological and immunohistochemical analyses were performed on samples derived from mouse tumor grafts and all analyzed parameters were blindly evaluated by the same pathologist (MP). In details, 4 µm-thick tissue sections were obtained from formalin-fixed paraffin-embedded tissue samples and representative tumor areas were selected on H&E-stained slides for immunohistochemical (IHC) analysis. IHC was performed using primary rabbit polyclonal anti HIF1α antibody (Novus Biologicals). Antigen retrieval was performed with heat/EDTA in the Bond-Max automated immunostainer (Leica Biosystems), as previously described (Pizzi et al., 2017).

### Quantification and statistical analysis

Data were analyzed and presented as mean ± standard deviation (SD) or standard error of the mean (SEM) in all figures. Pairs of data groups were analyzed using paired and unpaired two-tailed Student’s *t* tests. In the case of more than two groups, one-way analysis of variance (ANOVA) followed by Bonferroni post-hoc test was applied. Statistical significance was determined using Origin® 8 (OriginLab, Northampton, MA). Results with a p value lower than 0.05 were considered significant; ***p < 0.001, **p < 0.01, *p < 0.05 compared to controls. Each experiment was repeated at least three times.

## MATERIALS AVAILABILITY

Reagents generated in this study will be made available on request, but we may require a completed Materials Transfer Agreement.

## DATA AND CODE AVAILABILITY

Data are available upon request.

## KEY RESOURCES TABLE

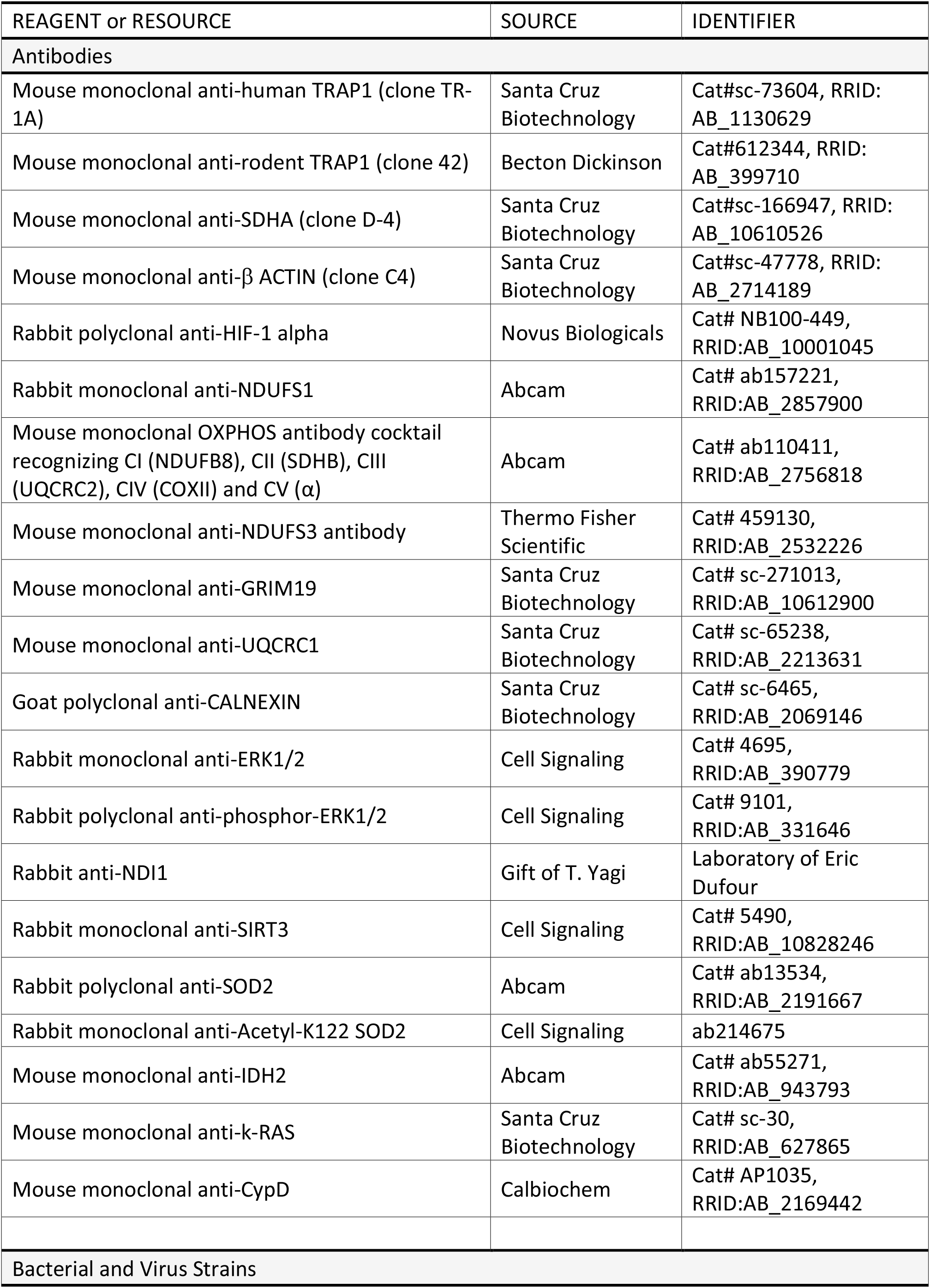
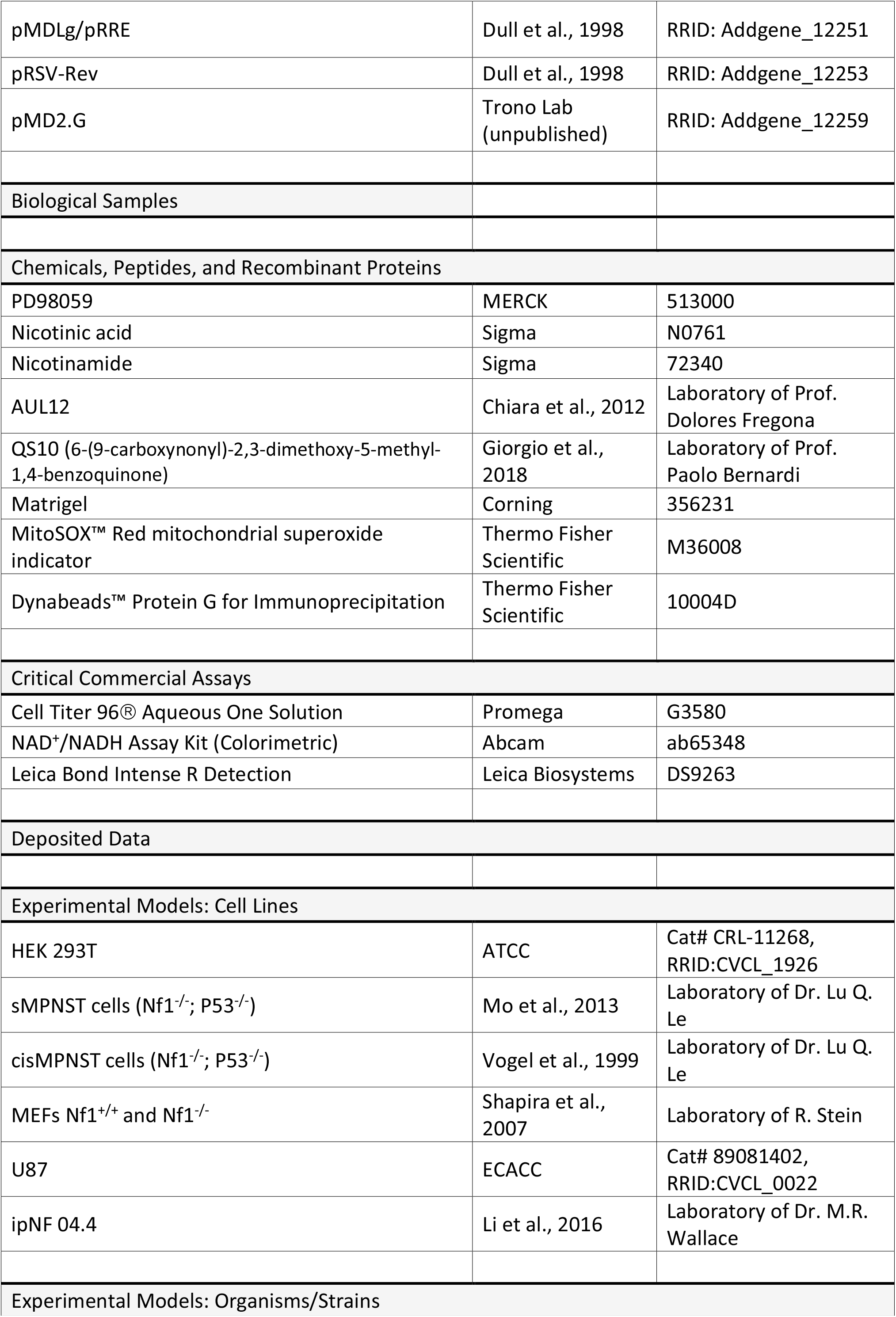
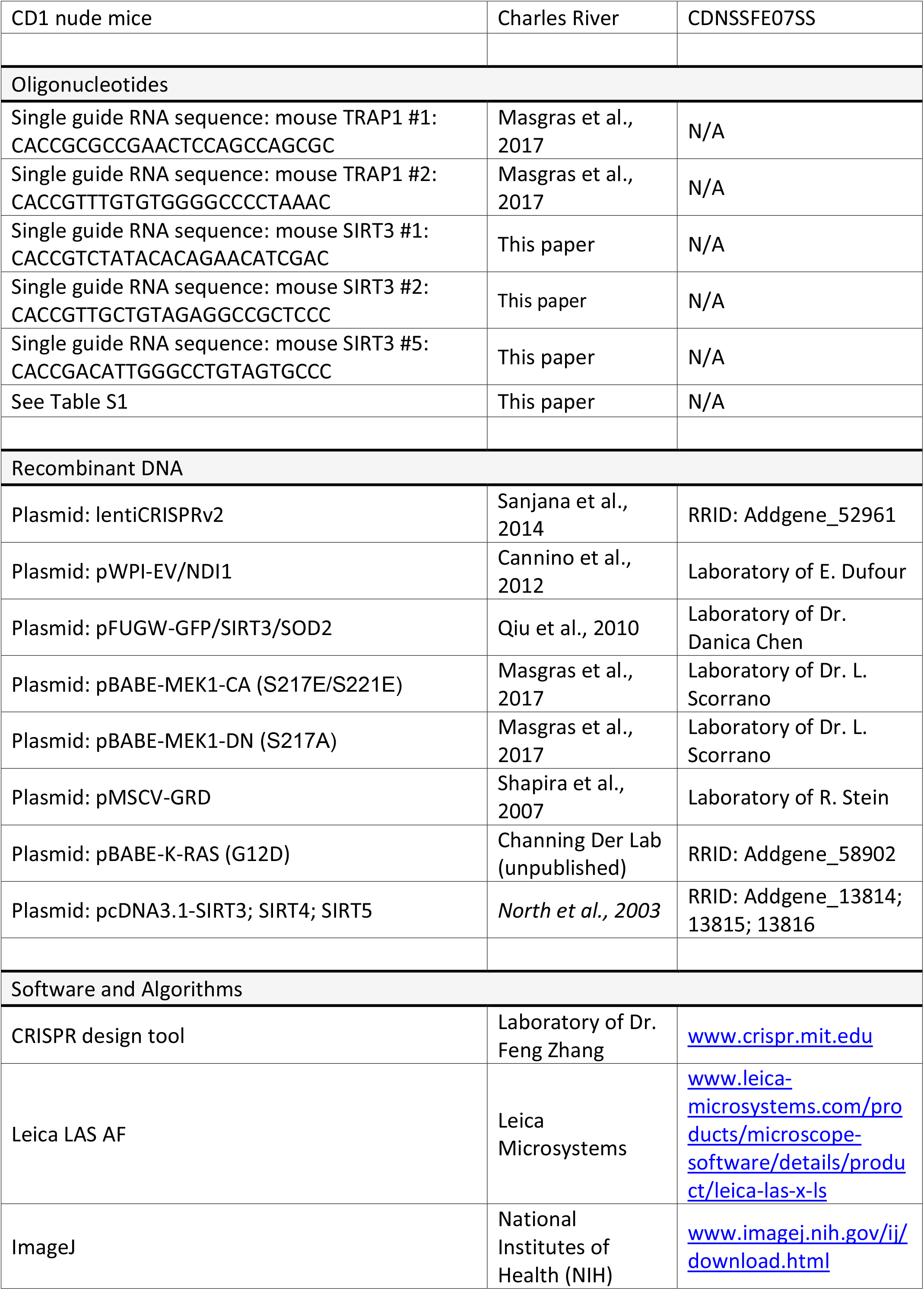
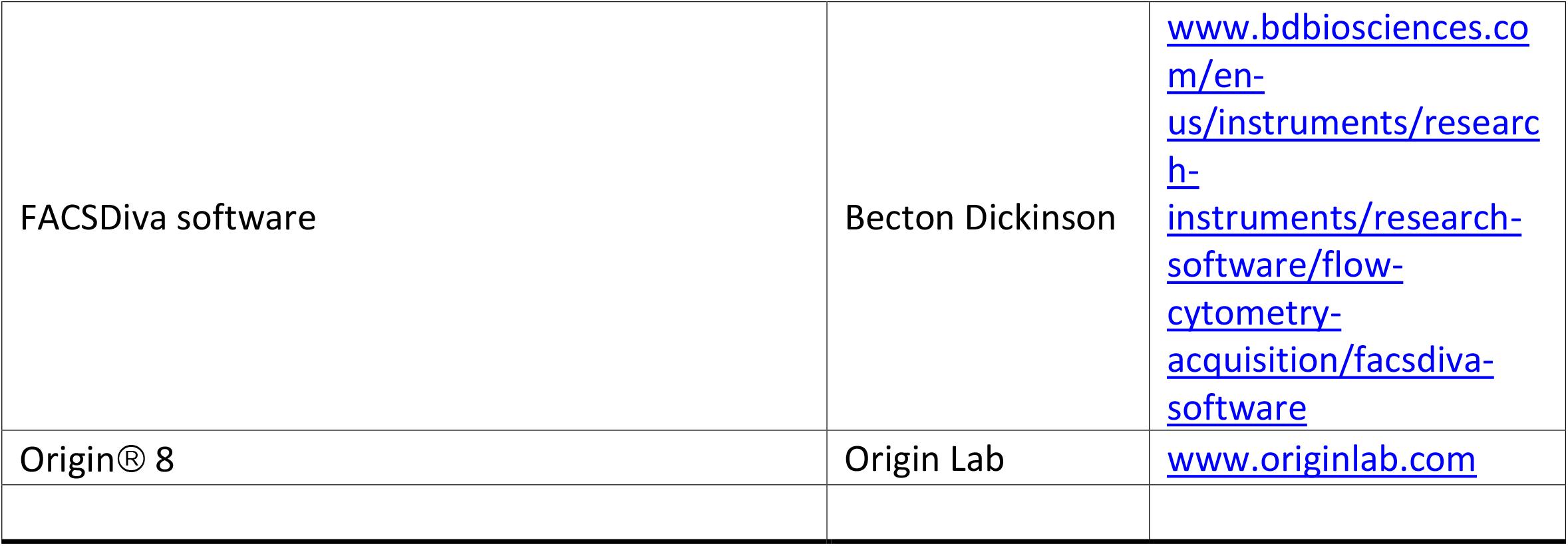

