## Supplemental information contains 6 supplemental figures and 1 supplemental table for "Tumor growth of neurofibromin-deficient cells is driven by decreased respiration and hampered by NAD^+^ and SIRT3"

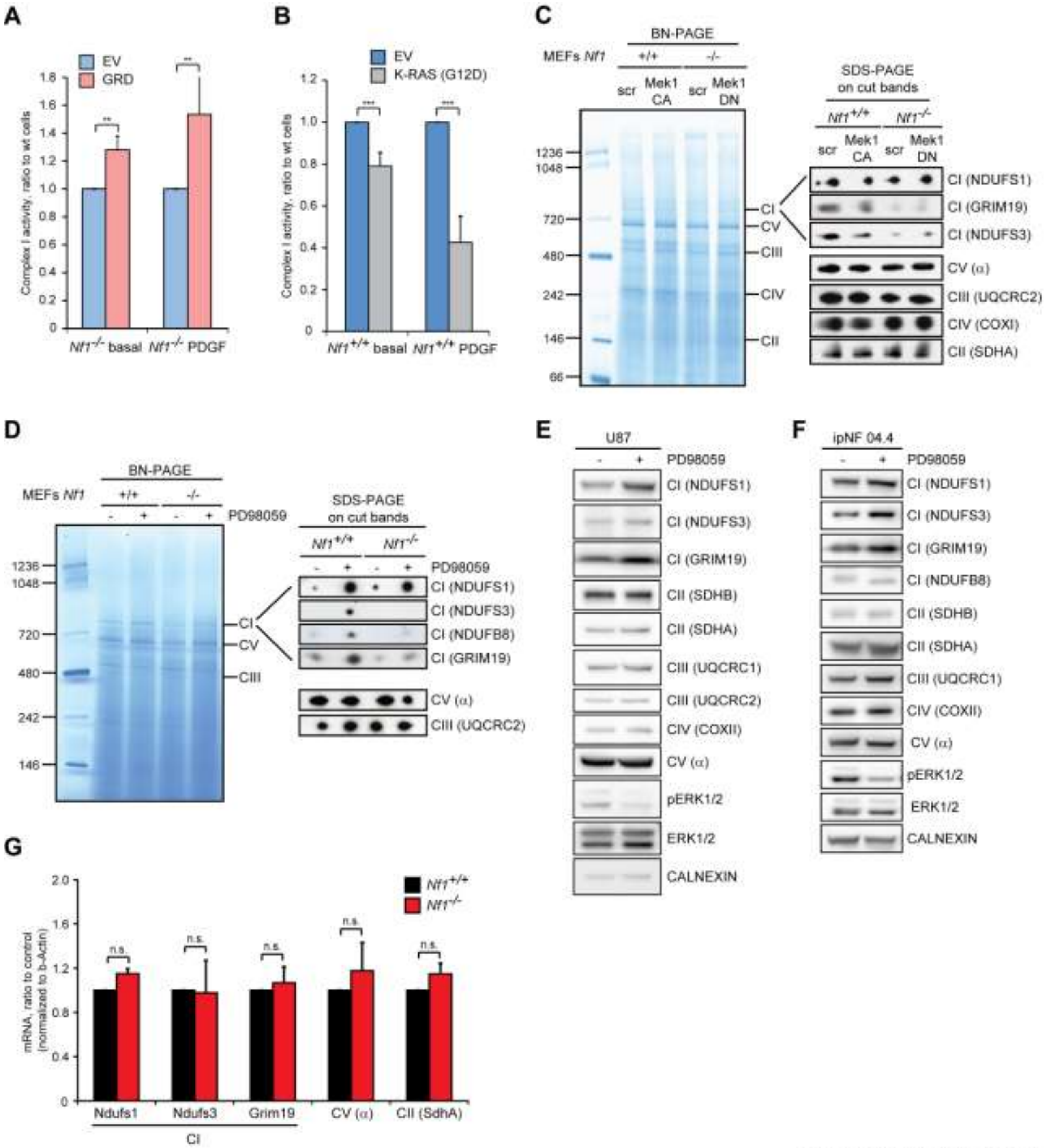

Supplementary Figure 1

**Supplementary Figure 1. Related to Figure 1. Protein levels and enzymatic activity of respiratory complex I inversely correlate with induction of Ras-MEK-ERK signaling.** Related to Figure 1.

(A and B) Spectrophotometric analysis of the NADH dehydrogenase activity of complex I (CI) in control (empty vector, EV) and GRD (A) or K-RAS (G12D) (B) expressing Nf1$^{-/-}$ and Nf1$^{+/+}$ MEFs, respectively.

(C and D) BN-PAGE of respiratory complexes performed on digitonized mitochondria from Nf1$^{+/+}$ and Nf1$^{-/-}$ MEFs. The MEK-ERK pathway is modulated by expression of the constitutively-active or of the dominant-negative form of the upstream MEK1 kinase (MEK1-CA and MEK1-DN, respectively; C) or by treatment with the MEK inhibitor PD98059 (40 μM, 3 days; D).

(E and F) WB analysis of OXPHOS proteins upon modulation of ERK activity by PD98059 treatment (40 μM, 3 days) in human U87 glioblastoma cells (E) and ipNF 04.4 plexiform neurofibroma cells (F). pERK1/2 indicates phosphorylated, active ERK1/2. Calnexin was used as a loading control. NDUFS1, GRIM19, NDUFS3 and NDUFB8 were used as complex I markers; subunits α, UQCRC1/C2, COXI and SDHA/B as complex V, complex III, complex IV and complex II markers, respectively.

(G) RT-PCR on mRNA levels of complex I, II and V subunits in Nf1$^{+/+}$ and Nf1$^{-/-}$ cells. Values were normalized for beta-actin mRNA expression.

Data are reported as mean ± SD values (n ≥ 3); ***:p<0.001; **: p<0.01; *: p<0.05 with a Student's *t* test analysis.

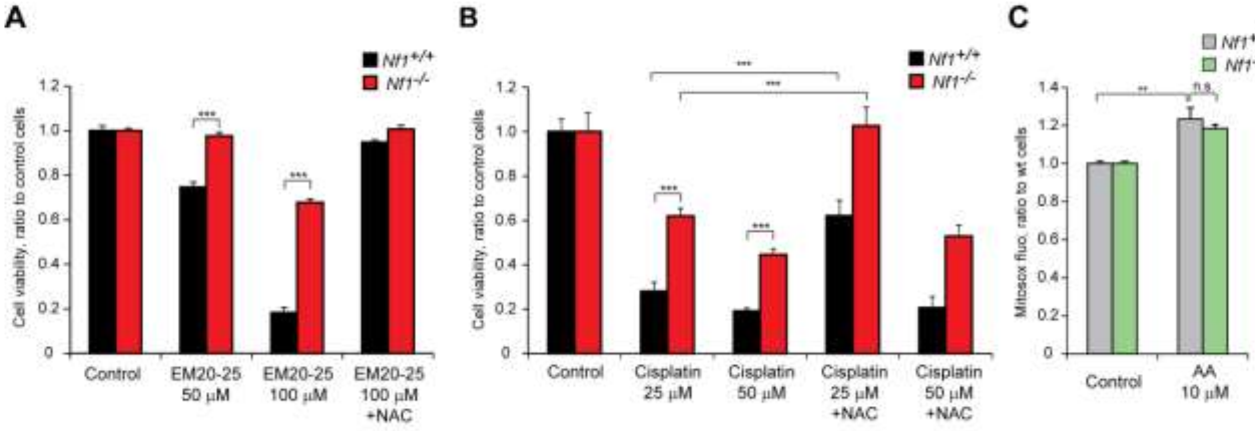


**Supplementary Figure 2**

**Supplementary Figure 2. Related to Figure 2. Neurofibromin loss decreases sensitivity to drugs targeting mitochondrial complex I.** Related to Figure 2.

(A and B) Cell viability following exposure to EM20-25 (50/100 μM, 18h; A) or cisplatin (25/50 μM, 24h; B) with or without NAC (500 μM) was assessed as in Figure 2C.

(C) Analysis of mitochondrial ROS levels by MitoSOX staining in control and antimycin A (AA)-treated (10 μM, 30 minutes). Experiments were carried out on Nf1$^{+/+}$ and Nf1$^{-/-}$ MEFs.

Data are reported as mean ± SD values (n ≥ 3); ***: p<0.001; **: p<0.01 and *: p<0.05 with a Student's *t* test analysis.

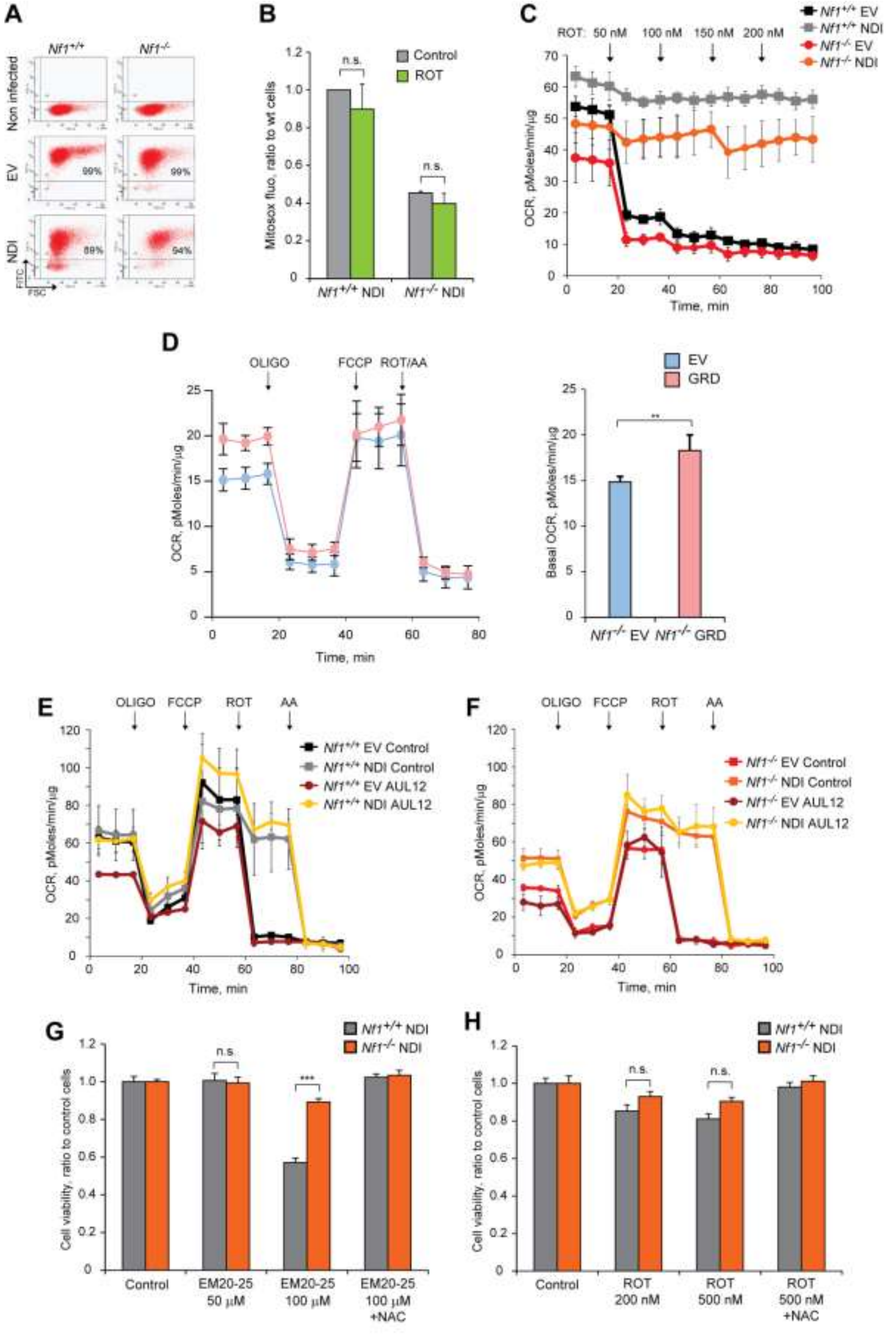


**Supplementary Figure 3**

**Supplementary Figure 3. Related to Figure 3. Effect of the alternative NADH dehydrogenase NDI1 on oxygen consumption rate (OCR), ROS and viability of cells.** Related to Figure 3.

(A) Expression of pWPI empty vector or pWPI-NDI1 (EV and NDI, respectively) fused with GFP was revealed by assessing green fluorescence in cytofluorimetric inspections.

(B) Analysis of mitochondrial ROS levels by MitoSOX staining in control and rotenone-treated (200 nM, 1 hour) NDI1-expressing cells.

(C) Representative OCR traces of cells harboring either pWPI empty vector (EV) or pWPI-NDI1 (NDI). The complex I inhibitor rotenone (ROT, 50-200 nM) was added where indicated.

(D) Representative OCR traces (left) and quantification of basal OCR values (right) in control and GRD-expressing $Nf1^{-/-}$ MEFs. The ATP synthase inhibitor oligomycin (0.8 μM), the proton uncoupler carbonyl cyanide-4-(trifluoromethoxy)phenylhydrazone (FCCP, 1 μM) and the respiratory complex I and III inhibitors rotenone (0.5 μM) and antimycin A (1 μM), respectively, were added where indicated.

(E and F) Representative OCR traces in control and AUL12-treated (4 μM, 45 minutes) $Nf1^{+/+}$ and $Nf1^{-/-}$ MEFs (D and E, respectively). The ATP synthase inhibitor oligomycin (0.8 μM), the proton uncoupler carbonyl cyanide-4-(trifluoromethoxy)phenylhydrazone (FCCP, 1 μM) and the respiratory complex I and III inhibitors rotenone (0.5 μM) and antimycin A (1 μM), respectively, were added where indicated.

(G and H) Analysis of cell viability following exposure of NDI-expressing MEFs to EM20-25 (50/100 μM, 18h; G) or rotenone (200/500 nM, 18h; H) in the presence or absence of NAC (500 μM).

Data are reported as mean ± SD values (n ≥ 3); ***: p<0.001; **: p<0.01 and *: p<0.05 with a Student's *t* test analysis. All experiments in the Figure were carried out on $Nf1^{+/+}$ and $Nf1^{-/-}$ MEFs.

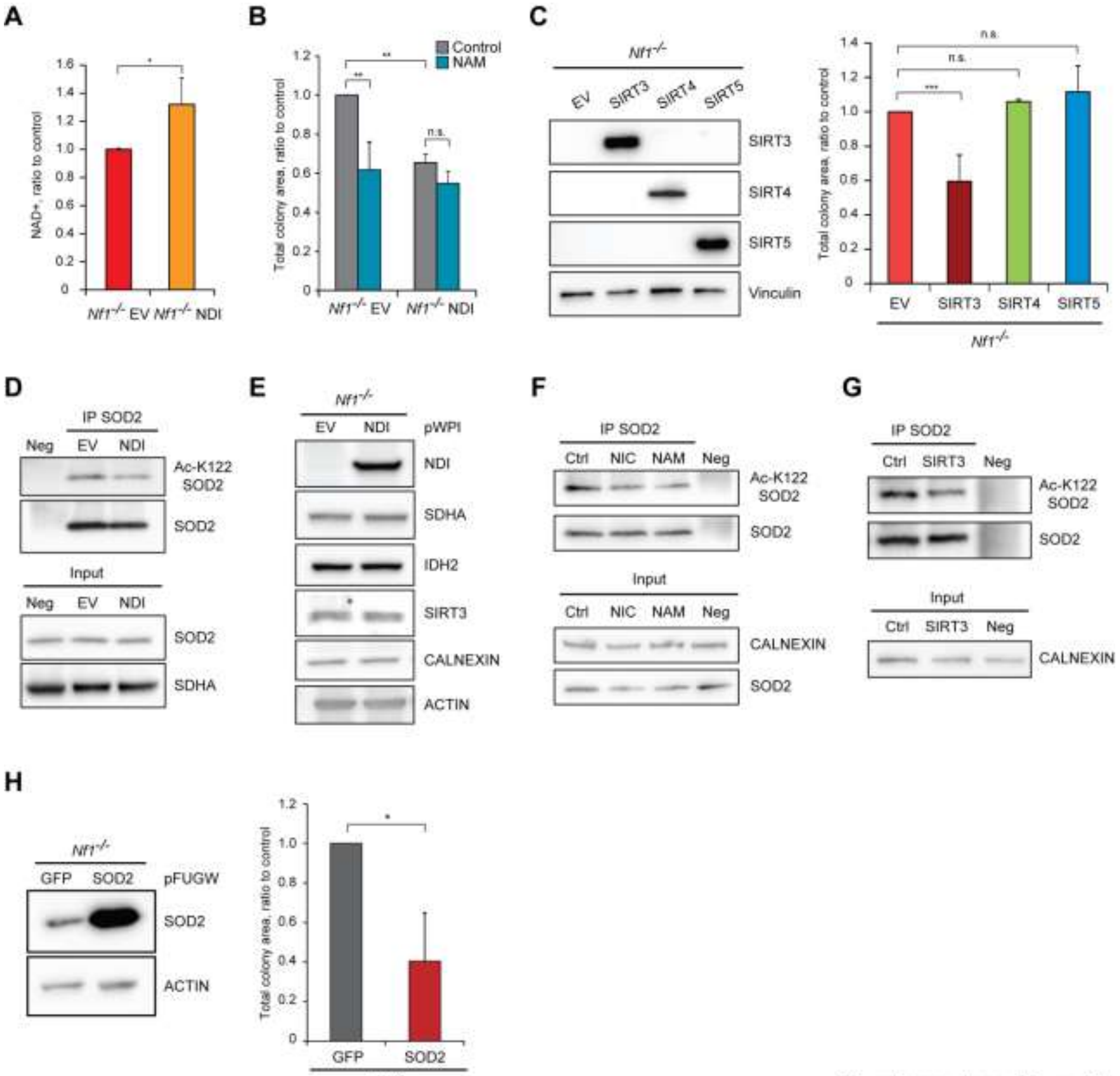


Supplementary Figure 4

**Supplementary Figure 4. Related to Figure 4. Effects of NDI1 and SOD2 expression on cells lacking neurofibromin.** Related to Figure 4.

(A) Measurement of $NAD^+$ level following NDI expression.

(B) Effect of nicotinamide (NAM) treatment on soft agar growth of Nf1$^{-/-}$ cells.

(C) Matrigel-embedded 3D colony growth (right) of Nf1$^{-/-}$ cells following SIRT3, SIRT4 or SIRT5 overexpression (left). Vinculin was used as loading control.

(D, F and G) SOD2 immunoprecipitation to assess acetylation of the SIRT3 target lysine 122 (Ac-K122 SOD2) following NDI expression (D), NIC/NAM treatment (F) and SIRT3 overexpression (G).

(E) WB analysis of the expression level of SIRT3 and of its targets succinate dehydrogenase subunit A (SDHA) and isocitrate dehydrogenase 2 (IDH2). Calnexin and actin were used as loading controls.

(H) SOD2 over-expression after transfection with pFUGW-SOD2 (left) decreased colony formation in soft agar (right). Negative controls were cells expressing pFUGW-GFP.

All experiments were carried out on Nf1$^{-/-}$ MEFs. NDI: cells expressing the pWPI-NDI1 construct; EV: cells expressing the pWPI empty vector.
Data are reported as mean ± SD values (n ≥ 3); ***: p<0.001; **: p<0.01 and *: p<0.05 with a Student's *t* test analysis.

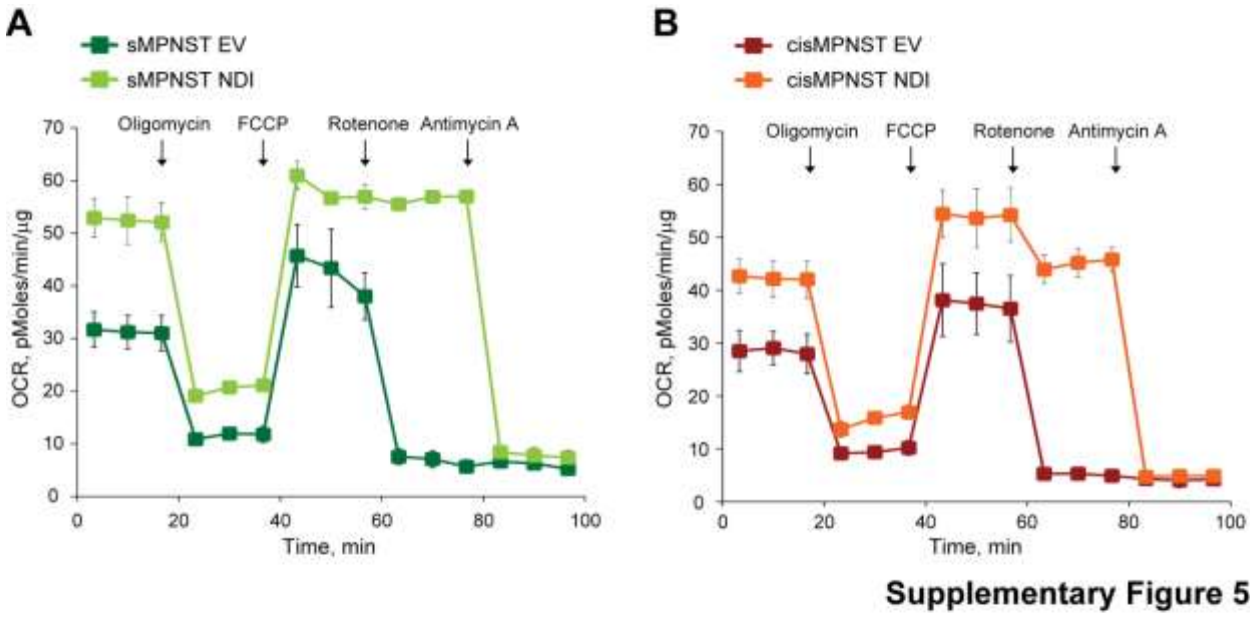


**Supplementary Figure 5. Related to Figure 5. Effect of NDI1 expression on bioenergetics of MPNST cells.** Related to Figure 5.
(A and B) Representative OCR traces of sMPNST (A) and cisMPNST (B) cell models. NDI: cells expressing the pWPI-NDI1 construct; EV: cells expressing the pWPI empty vector. The ATP synthase inhibitor oligomycin (0.8 µM), the proton uncoupler carbonyl cyanide-4-(trifluoromethoxy)phenylhydrazone (FCCP, 1 µM) and the respiratory complex I and III inhibitors rotenone (0.5 µM) and antimycin A (1 µM), respectively, were added where indicated.
Data are reported as mean ± SD values (n ≥ 3); ***: p<0.001; **: p<0.01 and *: p<0.05 with a Student's *t* test analysis.

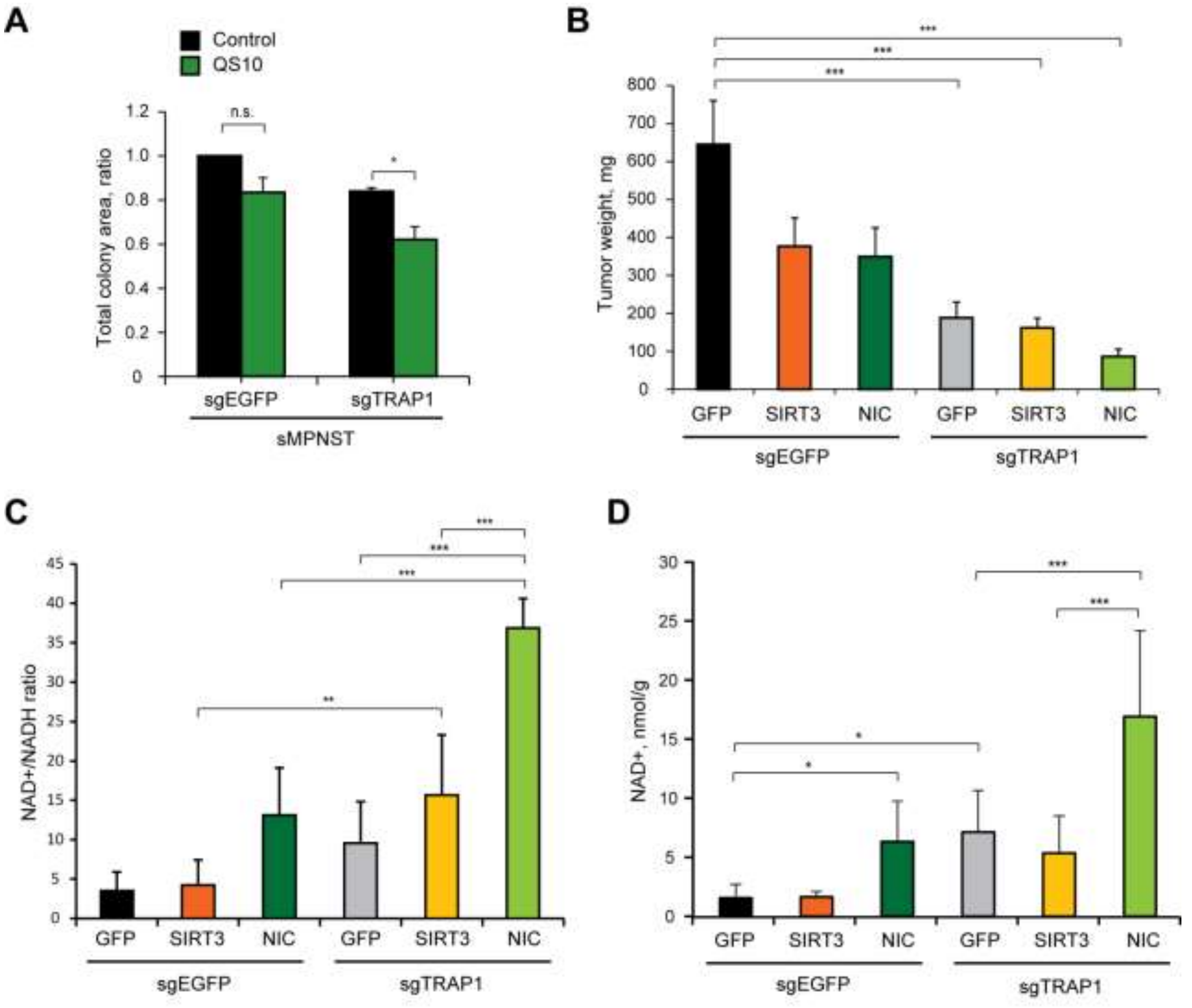


**Supplementary Figure 6**

**Supplementary Figure 6. Related to Figure 6. Anti-neoplastic effects of SIRT3 and NIC and of TRAP1 ablation.** Related to Figure 6.

(A) Effect of QS10 treatment (10 µM, 3 days) on colony growth in Matrigel of control and TRAP1 knock-out sMPNST cells.

(B, C and D) Measurement of tumor weight (mg; B), $NAD^+$/NADH ratio (C) and NAD+ (nmol/g; D) of tumor samples after xenograft injection in nude mice of control and TRAP1 knock-out sMPNST cells. Samples are from the experiments reported in Figure 6E.

All along the Figure, cells are labeled as in Figure 6A. In A, C and D data are reported as mean ± SD values (n ≥ 3). In B data are reported as mean ± SEM values (n ≥ 7). A Student's *t* test analysis was applied in A; a one-way ANOVA followed by Bonferroni post-test was performed in B, C and D. ***: p<0.001; **: p<0.01 and *: p<0.05.

**Table S1.** List of primers used for qPCR. Related to STAR Methods.

| Gene | Forward primer | Reverse primer |
|---|---|---|
| Actb | CCCCCTGAACCCTAAGGCCA | GGCTACGTACATGGCTGGGG |
| Sdha | CGGCTTTCACTTCTCTGTTGGTGA | AAAGGCCAAATGCAGCTCGCAA |
| Ndufs1 | TCTTCTGGGAGCAGATGGAGGT | ATGGGAGCACCAACATCACCA |
| Ndufs3 | ATGGCTTCGAGGGACATCCT | GGTTCAGCCACTACCCGCTT |
| Grim19 | CTACGGCCCCATCGACTACAAG | CCCCGATGCCCACAGCAAAC |
| ATP5a1 | GGCTGGTGATGTGTCCGCTT | TTTGGGCAGCAGATCCGACA |
